# Mechanical Rejuvenation of Mesenchymal Stem Cells from Aged Patients

**DOI:** 10.1101/2024.06.06.597781

**Authors:** Miles W. Massidda, Andrei Demkov, Aidan Sices, Muyoung Lee, Jason Lee, Tanya T. Paull, Jonghwan Kim, Aaron B. Baker

## Abstract

Mesenchymal stem cells (MSC) are an appealing therapeutic cell type for many diseases. However, patients with poor health or advanced age often have MSCs with poor regenerative properties. A major limiter of MSC therapies is cellular senescence, which is marked by limited proliferation capability, diminished multipotency, and reduced regenerative properties. In this work, we explored the ability of applied mechanical forces to reduce cellular senescence in MSCs. Our studies revealed that mechanical conditioning caused a lasting enhancement in proliferation, overall cell culture expansion potential, multipotency, and a reduction of senescence in MSCs from aged donors. Mechanistic studies suggested that these functional enhancements were mediated by oxidative stress and DNA damage repair signaling with mechanical load altering the expression of proteins of the sirtuin pathway, the DNA damage repair protein ATM, and antioxidant proteins. In addition, our results suggest a biophysical mechanism in which mechanical stretch leads to improved recognition of damaged DNA in the nucleus. Analysis of the cells through RNA-seq and ATAC-seq, demonstrated that mechanical loading alters the cell’s genetic landscape to cause broad shifts in transcriptomic patterns that related to senescence. Overall, our results demonstrate that mechanical conditioning can rejuvenate mesenchymal stem cells derived from aged patients and improve their potential as a therapeutic cell type.

**GRAPHICAL ABSTRACT:** 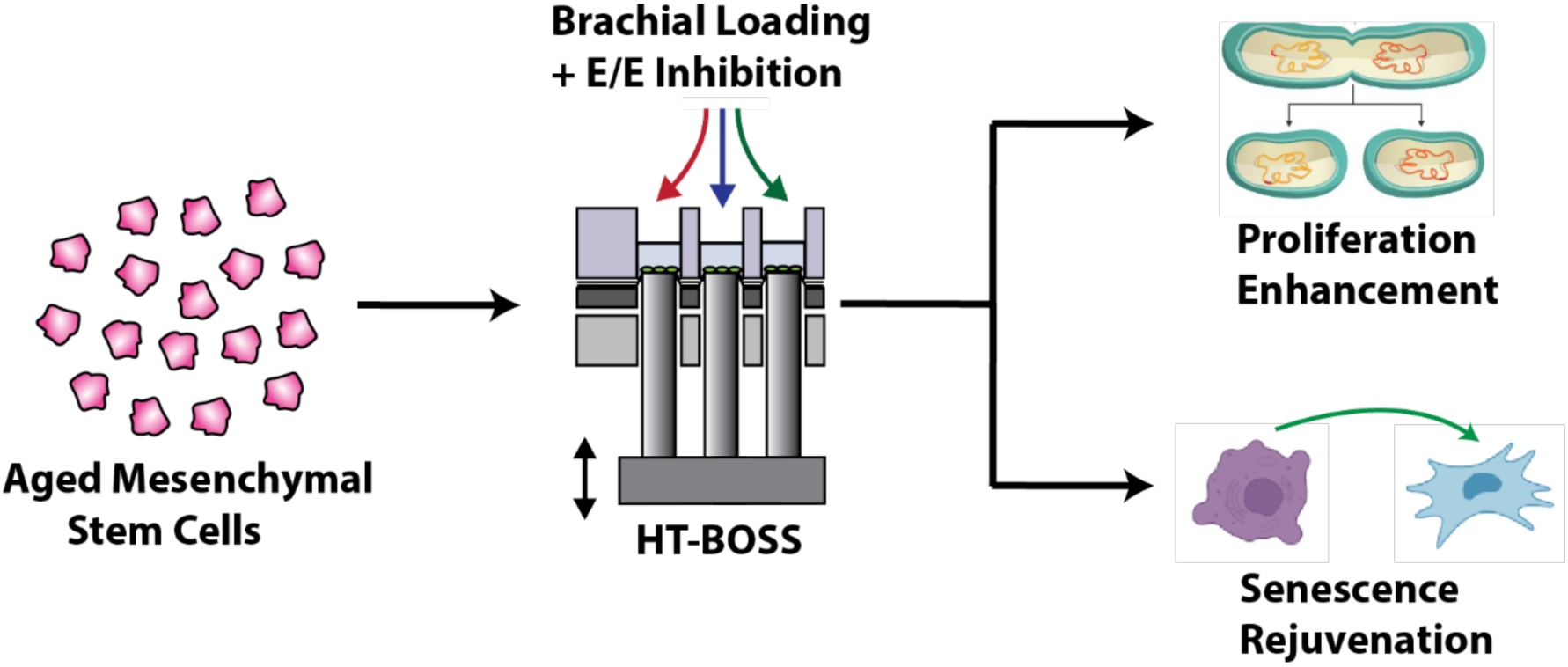

## INTRODUCTION

Bone marrow-derived mesenchymal stem cells (MSCs) are promising candidates for cell-based therapies for many diseases. MSCs are readily available in adult patients, do not require genetic modification for multipotency, and secrete bioactive factors for immunomodulation and angiogenesis.^1,2^ However, current MSC therapies have several key limitations, including suboptimal regenerative phenotype,^3^ limited potential for *in vitro* expansion and poor efficacy in aged or diabetic patients.^4^ A major key limiter of the regenerative function of MSCs is cellular senescence, which occurs more rapidly in MSCs from aged patients and during cell culture expansion.^5^ Senescence limits the efficacy of autologous MSC therapies in patients who are elderly^6,7^ or have co-morbidities like diabetes.^8,9^ Studies have also used MSCs derived from young, allogenic donors to bypass the hurdles poised by senescence in autologous therapies, but the long-term ability of allogenic MSCs to enhance tissue regeneration is limited by an immune response brought on by the transition of the cells from an immunoprivileged to immunogenic state following implantation in the patient.^10,11^ During culture expansion, MSCs experience a loss of differentiation potential and self-renewal capacity,^12^ shortening of telomeres and chromosomal instability,^13^ metabolic dysfunction,^14^ and loss of immunomodulatory functions.^15^ Thus, both allogenic and autologous MSCs are limited by cellular senescence due to cell culture expansion.

While the functional characteristics of MSC senescence are well described,^5,16,17^ the molecular mechanisms that contribute to the development of senescence are less well defined and often multifactorial. Senescent MSCs have limited proliferation capability,^18^ diminished multipotency,^19^ and reduced regenerative properties.^14^ Reactive oxygen species (ROS) byproducts are produced as MSCs age and replicate, causing stress-induced senescence, in which defective antioxidant mechanisms allow for ROS to build up in the mitochondria and lead to metabolic dysfunction and loss of proliferative capability.^20–22^ Oxidative stress and other stress stimuli can cause DNA damage, and dysregulated DNA damage repair pathways in aged MSCs can lead to apoptosis or the induction of senescence.^23,24^ Prior work has supported that targeting mechanisms of reducing oxidative stress can reduce senescence in MSCs.^14,25–27^

In this work, we performed a detailed and extensive analysis of how mechanical conditioning can be used to rejuvenate senescent MSCs and improve expansion in culture while maintaining their multipotency. Recent studies from our laboratory demonstrated that a physiological waveform of mechanical strain improved regenerative capacity of MSCs from young donors in the context of revascularization to ischemia and angiogenesis.^28^ This work demonstrated that mechanical loading with a physiological waveform (brachial waveform) at 7.5% mechanical strain in combination with a EGFR/ErbB2-4 (E/E) inhibitor leads to enhanced MSC production of soluble HGF, increases endothelial cell and pericyte markers, and pericyte-like functionality in MSCs.^28^ Using a high throughput biaxial oscillatory stretch system (HT-BOSS) developed in our laboratory, ^29–31^ we explore the effect of these loading conditions on senescence in MSCs derived from aged donors (68-92 years old). We found that mechanical conditioning led to a lasting enhancement of proliferative function, expansion capacity, multipotency, and reduction of senescence in MSCs from aged donors. Mechanical load and E/E inhibitor treatment was also found to alter the expression of proteins related to oxidative stress regulation and DNA damage repair pathways, suggesting that our mechanical and pharmaceutical treatments may rejuvenate aged MSCs through the activation of these pathways. A mechanistic evaluation of proteomic signaling in these pathways after brachial loading and pharmacological inhibitor co-treatment revealed that the mechanical enhancement of proliferation relied upon oxidative stress signaling and the amplification of antioxidant machinery. Our studies support that mechanical conditioning reduces senescent cell phenotypes and protects the cells from advancement of senescence through the activation of DNA damage repair signaling mediated by the ATM kinase protein. Lastly, integrated epigenetic and transcriptomic analyses revealed that mechanical loading of aged MSCs resulted in broad genome-wide changes in chromatin accessibility and gene expression that reverse the enrichment pattern of gene sets correlated with senescent cells.

## RESULTS

### Mechanical conditioning enhances proliferation in MSCs from aged donors

In previous studies, we identified specific mechanical loading regimes and pharmacological compounds to enhance the regenerative capacity of MSCs from young donors for treating peripheral ischemia.^28^ The specific conditions used a mechanical loading regime of 4 hours per day for 7 days of 7.5% maximal strain at 0.1 Hz frequency of loading using a physiological waveform similar to stretch of the brachial artery during the cardiac cycle (brachial loading).^32^ In addition, we identified that a EGFR/ErbB-2/4 (E/E) inhibitor as a co-treatment with loading led to a dramatic increase in MSCs that expressed the markers for both endothelial cells and pericytes, and observed that these cells had enhanced angiogenic activity with a tube formation assay.^28^ To test if mechanical loading could improve the proliferative function of MSCs from aged patients, we conditioned MSCs from three donors (68-92 years of age; one male and two female) with a brachial waveform at a maximal strain of 7.5% for 7 days for 4 hours per day in combination with 1 μM E/E inhibitor. After the treatments, cells were evaluated for DNA synthesis with a bromodeoxyuridine (BrdU) proliferation assay. Brachial loading was shown to increase the DNA synthesis of the cells in all three of the donors, and brachial + E/E inhibitor treatment increased DNA synthesis in two of three donor lines, relative to the static control **(Fig 1A)**. Additionally, treated cells from an 81-year-old female donor were collected and seeded into standard culture plates and passaged until they reached full senescence and failed to continue proliferating. Cells were counted at each passage, and brachial loaded cells were shown to maintain a significantly faster proliferative rate during passages 9-12, while static groups reached comparative growth arrest **(Fig 1B)**. To verify that biomechanically treated MSCs retained their phenotype after the treatment and during subsequent culture expansion, tested for the MSC phenotype of the cells using flow cytometry **(SI Fig. 1)**. Treated cells were evaluated for expression of proliferation-related proteins with western blotting **(Fig 1C, D)**. A Ponceau stain was run to confirm equal loading of cell lysate protein **(SI Fig. 2A)**. Results showed significant upregulation of cell-cycle regulators AKT (pan), p-AKT (Ser473), cyclin D1, and p-cyclin D1 (Thr286) with mechanical treatment alone (Brach).^33^

**Figure 1.**
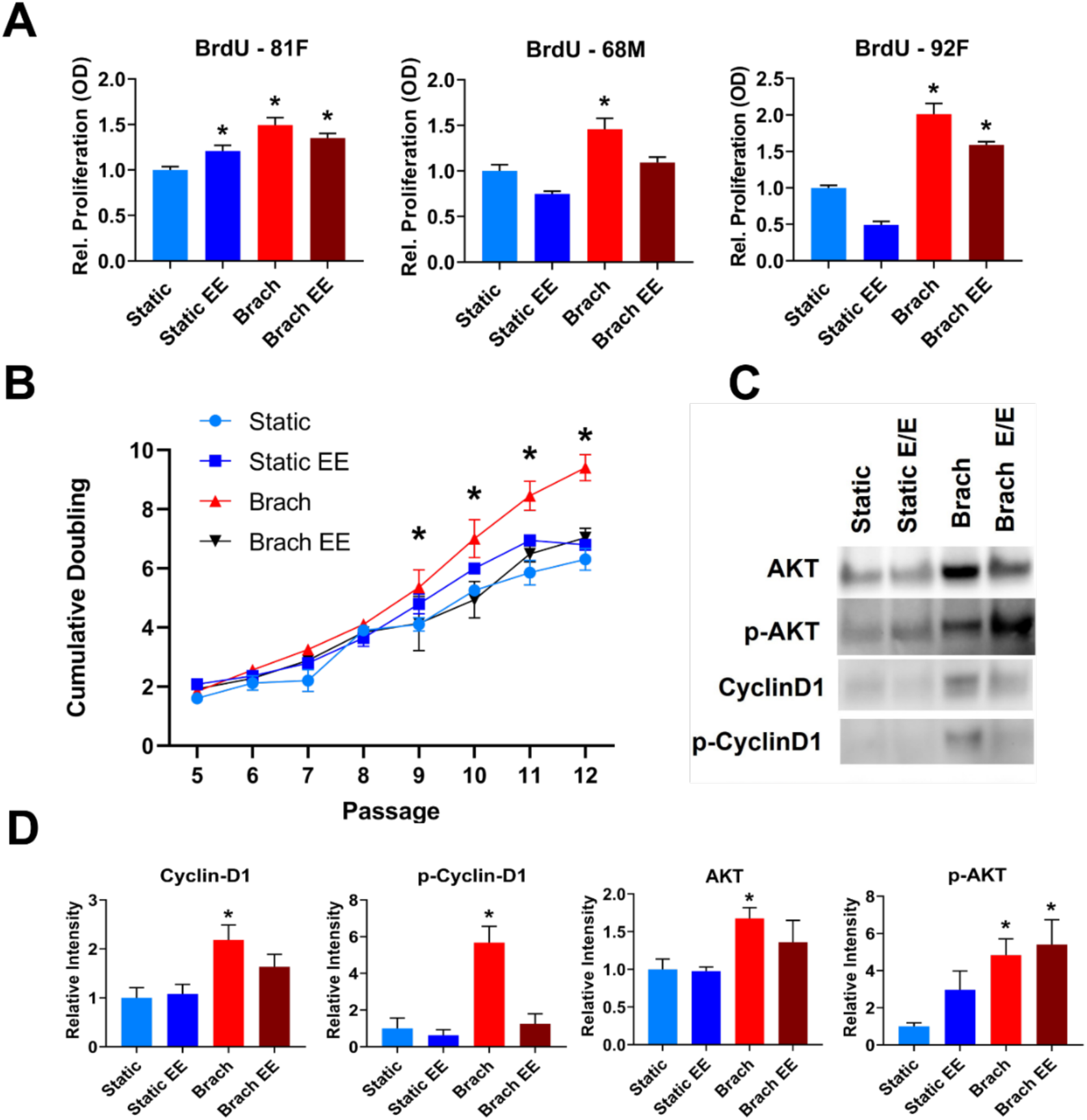
Mechanical and pharmaceutical treatments enhance proliferation of aged MSCs. MSCs derived from three aged donors were treated with brachial loading and E/E inhibitor and assessed for potential rejuvenation of proliferative ability. **(A)** Quantification of BrDU cell proliferation assay. \**p <* 0.05 vs. static control treatment. **(B)** Quantification of the cumulative doubling of the treated cells from an 81-year-old female donor during long-term cell culture expansion. *p < 0.05 vs. static control treatment. **(C)** Western blotting results and **(D)** quantification for proliferation-related protein expression after treatment of MSCs. *p < 0.05 vs. static control treatment.

**Figure 2.**
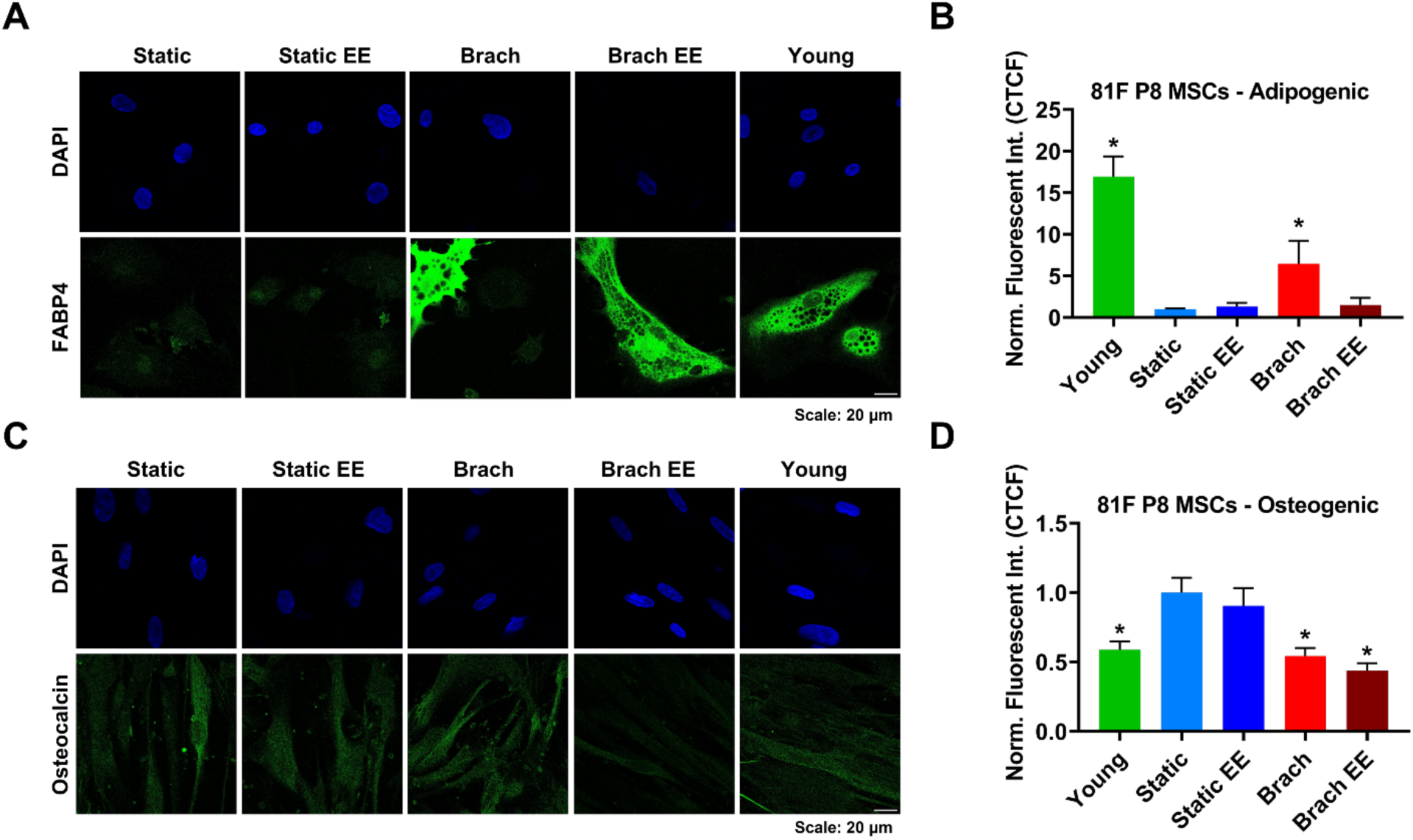
Mechanical conditioning enhances differentiative capability of aged MSCs. MSCs derived from one aged donor were treated with brachial loading and E/E inhibitor, expanded until P8, then differentiated into adipogenic and osteogenic lineages. **(A)** Adipogenic staining results and **(B)** quantification FABP4+ cells. *p < 0.05 vs. static control treatment. Scale bar = 20 µm. **(C)** Osteogenic staining results and **(D)** quantification of Osteocalcin+ cells. *p < 0.05 vs. static control treatment. Scale bar = 20 µm.

### Mechanical conditioning enhances differentiative capability of aged MSCs

Senescent MSCs are often shown to have a reduced capacity to differentiate into other cell phenotypes, which limits their clinical utility.^19^ To test if mechanical conditioning could improve the multipotency of aged MSCs, we repeated brachial loading and E/E inhibitor treatments on P5 MSCs from an 81-year-old female donor. Following the treatments, we expanded the MSCs in culture until P8, then evaluated for adipogenic and osteogenic differentiative potential with the hMSC functional identification kit. A young MSC line from a 24-year-old female donor was also expanded until P8 and included as a control. The rationale for using cells at P8 is that senescent MSCs will have a significant loss in their ability to undergo functional tri-linage differentiation, and studies have shown that an almost complete loss in multipotency occurs around P9/P10 for cells derived from old donors (60+ years).^34^ Thus, we tested just prior to the the point of complete induction of senescence to determine if our treatments can improve impaired multipotency. Following differentiation, cells were fixed and stained with FABP4 to evaluate adipogenic differentiation and Osteocalcin to evaluate osteogenic differentiation (**Fig. 2**). Results showed a significant increase in adipogenic differentiated cells with brachial loading, in comparison to the static control (**Fig. 2A, B**). We observed a decrease in osteogenic differentiation with brachial loading, but this matched the trend of the young control line, suggesting that our treatments may cause the cells to adopt a phenotype more similar to cells from younger donors (**Fig. 2C, D**). Furthermore, other studies have shown that the expression of osteogenic genes including osteocalcin tend to increase during MSC aging and *ex vivo* expansion, suggesting that our treatment may be inhibiting or reversing this process.^35^

### Mechanical conditioning upregulates Sirtuin expression in aged MSCs

The sirtuin family of proteins are nicotinamide dinucleotide (NAD+) dependent enzyme deacylases that regulate many cellular processes, including DNA repair, differentiation, metabolism, inflammation, and aging.^36^ Several members of the sirtuin family have also been implicated as major regulators of multiple senescence pathways.^40,64^ To investigate the effects of mechanical and pharmaceutical conditioning on sirtuin expression, we treated MSCs from an 81-year-old donor with brachial loading and E/E inhibition and performed western blotting for several proteins in the sirtuin family. Ponceau stain was run to confirm equal loading of cell lysate protein **(SI Fig. 2A)**. Results showed significant upregulation of SIRT1, p-SIRT1, a SIRT2 isoform, and SIRT6 with brachial loading, and upregulation of p-SIRT1, SIRT6, and SIRT7 with brachial + E/E treatment, in comparison to the static control **(Fig 3. A, B)**. This broad activation of sirtuin expression suggests that the mechanical and pharmaceutical treatments are altering signaling in age or senescence-related pathways, possibly to rejuvenate the proliferative and multipotent functions of the MSCs.

**Figure 3.**
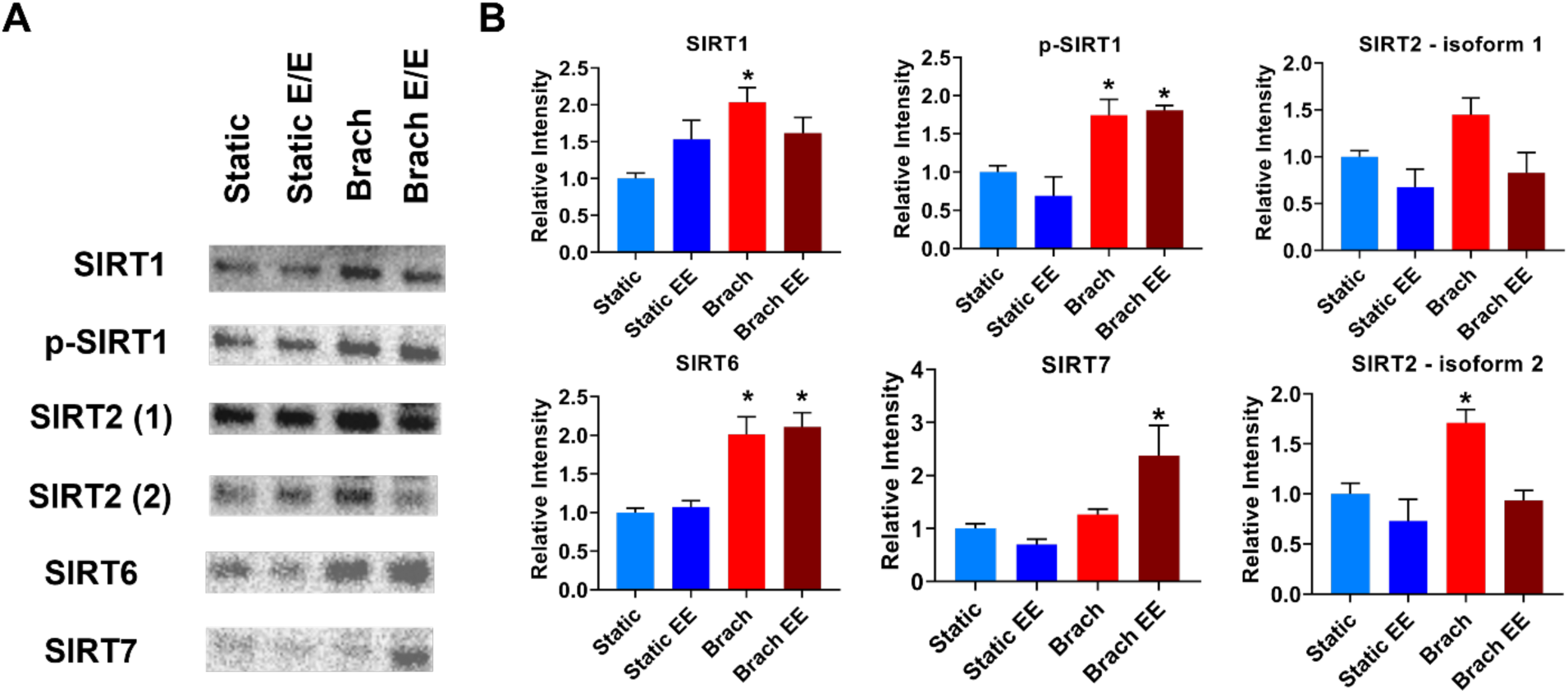
Mechanical and pharmaceutical treatments upregulate Sirtuin expression in aged MSCs. MSCs derived from one aged donor were treated with brachial loading and E/E inhibitor and assessed for expression of several proteins in the Sirtuin family. **A)** Western blotting results and **(B)** quantification of Sirtuin protein expression after treatment of MSCs. *p < 0.05 vs. static control treatment.

### Mechanical conditioning enhances oxidative stress management in aged MSCs

We next focused on the role of mechanical conditioning on key mechanisms in the development and progression of senescence, oxidative stress signaling and DNA damage repair. The effective management of oxidative stress and repair of DNA damage are critically important mechanisms for preventing senescence in MSCs.^24,37,38^ To test if mechanical loading could alter these mechanisms, we repeated mechanical and pharmaceutical treatments on MSCs from an aged donor (age 81) and evaluated for protein expression with western blotting in pathways related to oxidative stress (**Fig. 4A. B**). A Ponceau stain was included to confirm equal loading of cell lysate protein **(SI Fig. 1B)**. Results showed significant upregulation of FOX0 transcription factors, which are critical mediators of the cellular response to ROS in MSCs.^39–41^ Brachial loading or brachial + E/E inhibitor treatment caused an upregulation of FOX01, p-FOX01, FOX03a, p-FOX03a, and p-FOX04. Additionally, the treatments caused significantly upregulated expression of the antioxidant enzyme SOD1.^42^ (**Fig. 4A, B**). Next, we conditioned the MSCs with brachial loading or static control treatment for 4 hours and performed staining for oxidative stress with the CellROX Deep Red Reagent (ThermoFisher). We included the Sytox Blue dye (ThermoFisher) to distinguish dead cells from viable cells and performed flow cytometry. Results showed that mechanically conditioned cells had a significantly lower expression of reactive oxygen species (ROS) in comparison to the static control (**Fig. 4C**). These results indicate that the brachial loading treatment is altering the cellular response to oxidative stress signaling and mitigating ROS buildup. Studies have shown that oxidative stress often precedes senescence, leading to lessened proliferation, accelerated growth arrest, and DNA damage-related genetic instability.^37,43^

**Figure 4.**
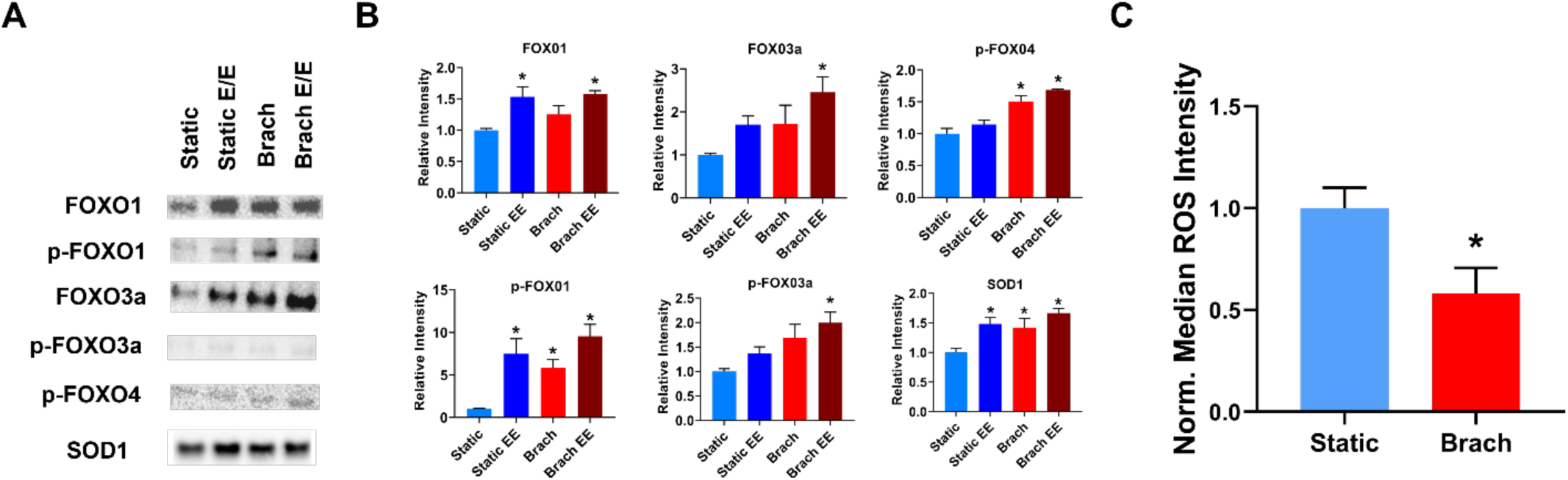
Mechanical conditioning enhances oxidative stress management in aged MSCs. MSCs derived from an 81-year-old female donor were treated with brachial loading and E/E inhibitor for 4 hours per day for 7 days and assessed for proteomic signaling in key pathways related to oxidative stress. **(A)** Western blotting results and **(B)** quantification of oxidative stress signaling protein expression after treatment of MSCs. *p < 0.05 vs. static control treatment. The MSCs were treated with brachial loading or static control treatments for 4 hours, and flow cytometry for the detection of reactive oxygen species (ROS) was performed. **(C)** Quantification of median ROS signal intensity. *p < 0.05 vs. static control treatment.

### Mechanical conditioning enhances the recognition and repair of DNA damage in aged MSCs

Studies suggest that long term culture of MSCs impairs the ATM-dependent recognition of DNA breaks, leading to the buildup of DNA damage and genetic instability.^44^ To investigate the effects of mechanical loading on DNA damage repair signaling, we repeated mechanical and pharmaceutical treatments on MSCs from an aged donor (age 81) and evaluated for protein expression with western blotting in pathways related to DNA damage repair (**Fig. 5**). A Ponceau stain was run to confirm equal loading of cell lysate protein **(SI Fig. 2B)**. Results showed significant upregulation of DNA-damage repair proteins KU80,^45^ ATM,^46^ and p-ATM^47^ with mechanical treatment (**Fig 5A,B**).

**Figure 5.**
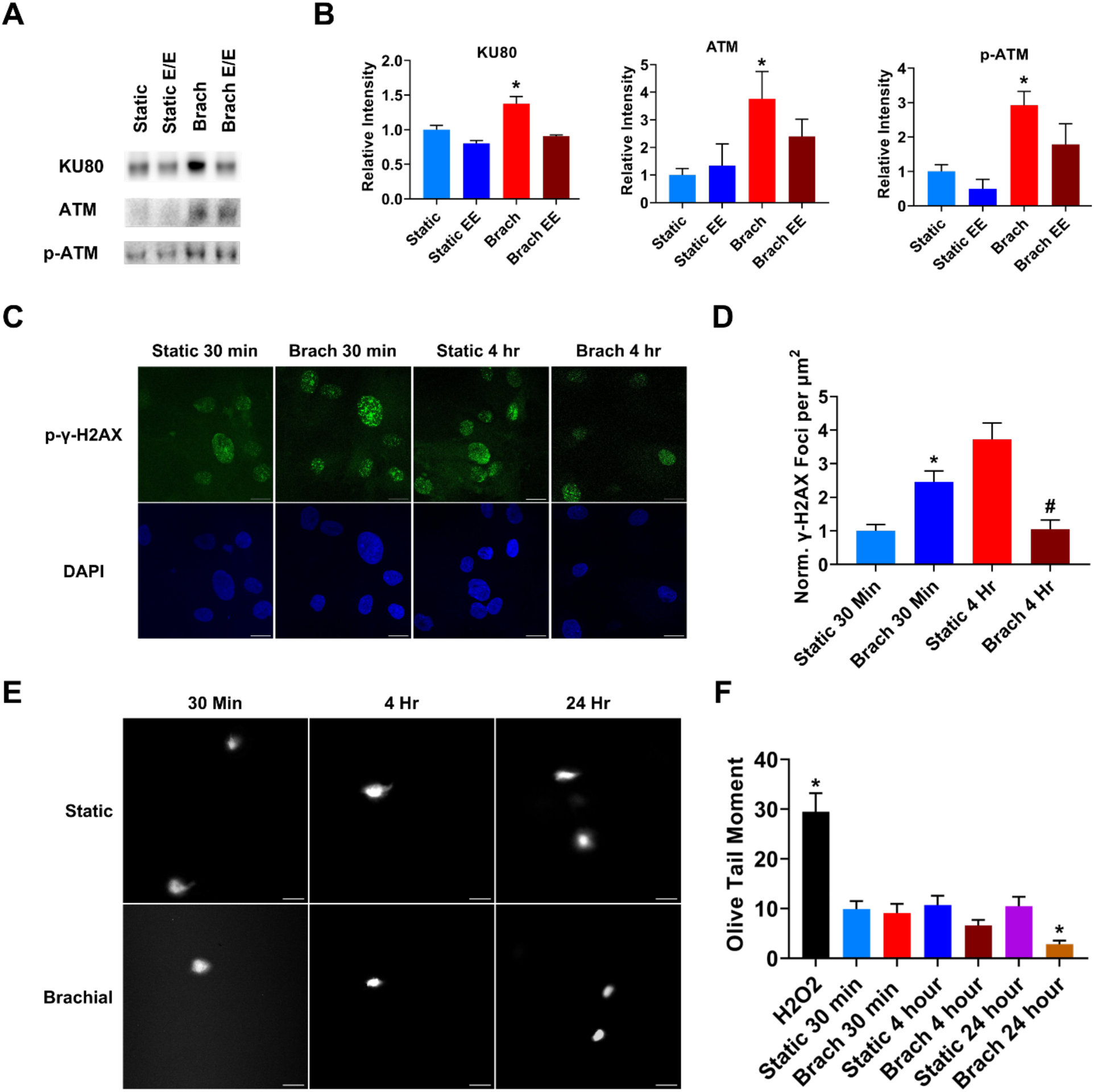
Mechanical conditioning enhances DNA damage recognition and repair in aged MSCs. MSCs from an 81-year-old female donor were treated with brachial loading and E/E inhibitor for 4 hours per day for 7 days and assessed for proteomic signaling in key DNA-damage repair pathways. **(A)** Western blotting results and **(B)** quantification of DNA damage-related protein expression after treatment of MSCs. *p < 0.05 vs. static control treatment. MSCs were treated with brachial loading or static control treatments for 30 minutes, or 4 hours. **(C)** Immunostaining of p-γ-H2AX histone foci and DAPI. Scale bar = 20 µm. **(D)** quantification of p-γ-H2AX histone foci expression. *p < 0.05 vs. 30-minute static control treatment. #p < 0.05 vs. 4-hour static control treatment. Brachial conditioning was repeated for 30 minutes, 4 hours, or 4 hours of treatment followed by 20 hours of static treatment. **(E)** Neutral comet assay images and (**F**) Quantification of DNA damage through calculation of the Olive Tail Moment (OTM) with the OpenComet plugin in FIJI. Scale bar = 50 µm.

To investigate the downstream effects of mechanically altered DNA damage repair signaling, we treated MSCs from the 81-year-old donor with brachial loading or static conditions for 30 minutes, and 4 hours. Next, we performed immunostaining for p-γ-H2AX DNA double-strand break (DSB) repair foci. The histone H2AX makes a critical contribution to genetic stability through signaling the recognition of DNA damage events and acting as a foundation for the assembly of repair machinery.^48^ Results showed that brachial treatment for 30 minutes resulted in a significantly higher count of p-γ-H2AX foci compared to the static control at the same time point. However, after 4 hours of brachial loading, p-γ-H2AX foci was significantly reduced compared to the 4-hour static control group (**Fig. 5C, D**). This result suggests several possible mechanisms, one could be that brachial loading induces DNA damage leading to enhanced DNA damage repair at a later time another is that the mechanical stretch is revealing occult DNA damage that is not recognized by the cell until the stretch acts to open the chromatin. To investigate these possibilities further, we performed a neutral comet assay to assess the total accumulation of DNA DSBs at each time point.^49^ This assay does not require the cell’s recognition mechanism to detect DSBs so it able to detect the double stranded breaks independent of the DNA damage repair mechanism. The comet assay revealed that DNA damage remained relatively constant after 30 minutes and 4 hours of brachial treatment, compared to the static control (**Fig. 5E, F**). These data suggest that brachial treatment could be revealing existing DNA DSBs, rather than inducing new DNA damage. Furthermore, we also mechanically conditioned the cells for 4 hours, then left the cells in static conditions for an additional 20 hours, following the methods of the typical 7-day mechanical treatment used throughout this study. We found that after 24 hours, the brachial-treated groups demonstrated significantly less DNA damage than the static control. Altogether, these results suggest that brachial loading rapidly reveals sites of existing DNA damage through p-γ-H2AX foci expression, then enhances the repair of this damage to restore genetic stability. Our group recently also created a computational model of a cell being stretched on a flexible membrane.^50^ This model supports that the loading conditions used would be sufficient to induce opening of chromatin without damage to the DNA.

We next analyzed the nuclear morphology of the mechanically treated MSCs with the Nuclear Irregularity Index (NII) plugin in FIJI.^51^ MSCs from a young donor (24 year old female) or an aged donor (81 year old female) were treated with brachial loading or static control treatment for 4 hours. Cells were immunostained for DAPI and processed with the NII plugin to characterize their nuclei as “Normal”, “Apoptotic”, “Senescent”, or “Irregular” **(SI Fig. 3**). Results demonstrate that brachial loading causes a significant increase in the frequency of cells with “normal” nuclei, as well as a significant decrease of cells with “irregular” nuclei **(SI Fig. 3B**).

To further investigate the mechanism by which mechanical loading enhances the recognition and repair of DNA damage, we repeated the mechanical treatment of aged MSCs in combination with several pharmaceutical inhibitors of nuclear mechanotransductive signaling. Previous studies have demonstrated that the dynamic tensile loading of MSCs results in altered chromatin condensation that persists long after the completion of mechanical conditioning.^52^ This phenomena, deemed “mechanical memory”, may provide rationale for the long-term functional benefits demonstrated in this study after mechanical conditioning of aged MSCs. Previous work established that mechanically-driven changes in MSC chromatin architecture are largely dependent on Piezo mechanosensitive calcium channel signaling.^52^ To investigate the role of mechanosensitive signaling in enhanced DNA damage repair, MSCs from one aged donor were treated with brachial loading or static control treatment for 4 hours and co-treated with 40 µM Importazole, a drug that blocks importin-β-dependent nuclear import,^53^ 2 µM Dooku 1, which blocks Yoda1-induced activation of the Piezo1 Ca^2+^ mechanosensitive ion channel,^54^ or GsMTx4, which directly inhibits Piezo1 channel activity.^55^ Following the treatments, MSCs were immunostained for p-γ-H2AX foci **(SI Fig. 4)**. Results demonstrate that enhanced repair of DNA DSB foci is not dependent on mechanosensitive Piezo1 activation **(SI Fig. 4B**). Interestingly, co-treatment with brachial loading and Importazole resulted in significantly more p-γ-H2AX foci than treatment with brachial loading alone. This result suggests that while brachial loading is able to mechanically reorganize damaged DNA to enable its repair, this mechanism may be partially dependent on importin-β nuclear transport of proteins such as transcription factors to mediate DNA damage signaling.^56,57^ Furthermore, GsMTx4 and Dooku1 treatment of static control cells caused significantly more p-γ-H2AX foci than static treatment alone, suggesting that Piezo1-mediated nuclear influx of Ca^2+^ may regulate DNA damage response in MSCs.^58^

### Alteration of ATM-mediated DNA damage repair and oxidative stress signaling reduces mechanical protection from senescence and proliferation enhancement

Next, we evaluated the role of oxidative stress signaling and ATM-mediated DNA damage repair in mechanically rejuvenating the proliferative function of aged MSCs and protecting them from senescence. MSCs were treated with 5 µM of KU55933 (an ATM kinase inhibitor, **ATMi**),^59^ 5 mM of N-acetyl-L cysteine (**NAC**), which has been shown to have strong antioxidant effects in MSCs,^60^ or control treatments. Cells were then mechanically conditioned and evaluated for DNA synthesis with a BrdU assay. Results showed significantly higher DNA synthesis under brachial loading and brachial + KU55933 treatment, but not under brachial + NAC treatment **(Fig. 6A).** These results suggest that brachial loading may rely on oxidative stress signaling and the amplification of antioxidant machinery to enhance the proliferative function of the cells. Next, we repeated the mechanical and pharmaceutical treatments and performed staining for β-galactosidase, a common biomarker of senescent cell phenotypes.^37,61,62^ Results showed a significant reduction in the percentage of β-galactosidase+ cells in NAC-treated groups and brachial-treated groups, but not KU55933-treated groups (**Fig. 6B, SI Fig. 5**). These results demonstrate the potential of mechanical conditioning to either protect the cells from advancing senescence, or rejuvenate senescent cells into a healthy phenotype, generating a lower population of senescent cell phenotypes following the treatment. The data also suggests that, specifically, the mechanical enhancement of ATM-mediated DNA damage repair signaling is critical for protecting aged MSCs from the progression of senescence.

**Figure 6.**
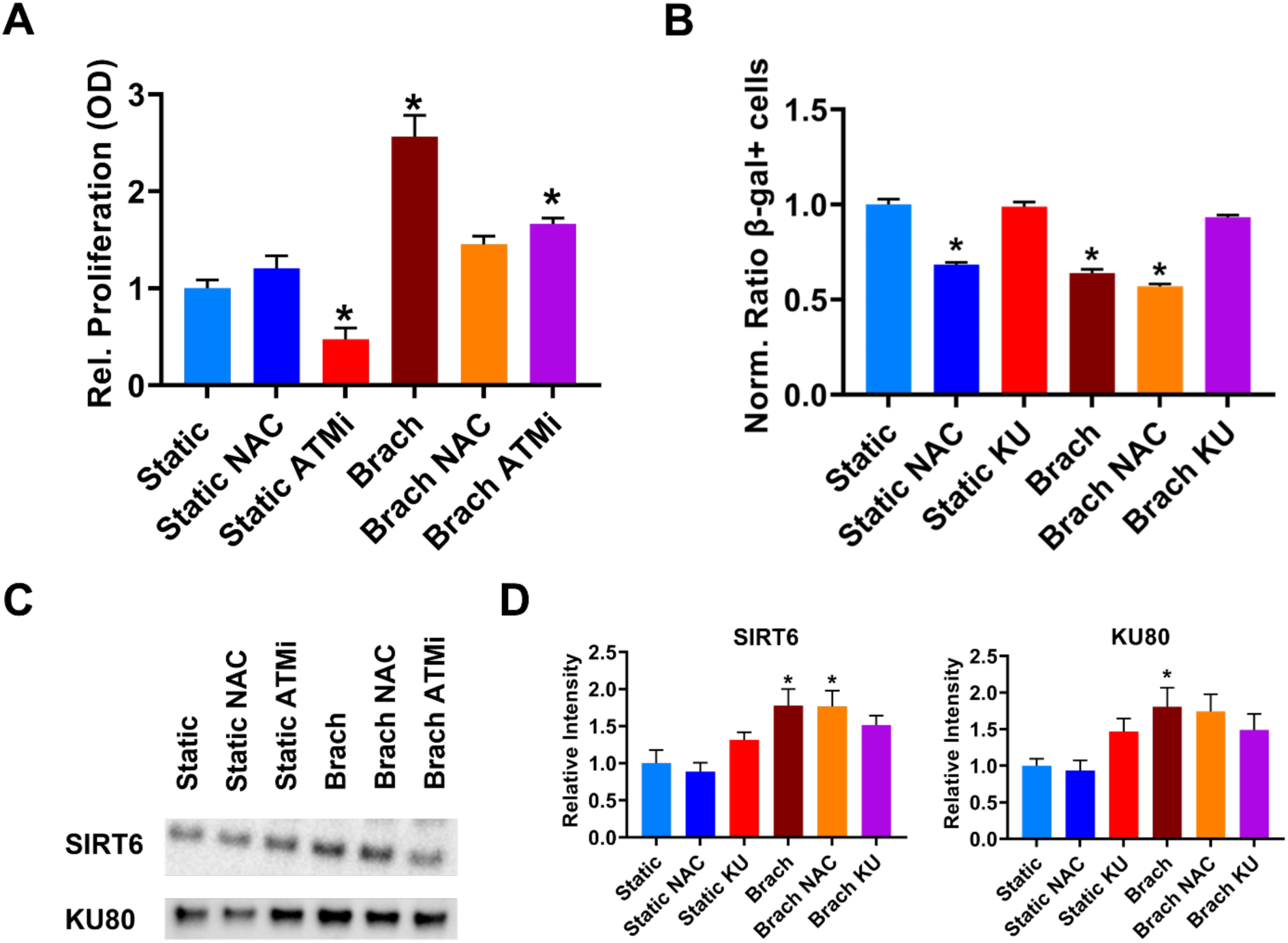
Alteration of ATM-mediated DNA damage repair and oxidative stress signaling reduces mechanical protection from senescence and proliferation enhancement. MSCs derived from one aged donor were co-treated with antioxidant N-Acetyl Cysteine (NAC), ATM inhibitor KU55933 (ATMi), and brachial loading. Following the treatments, MSCs were = assessed for proliferative function, senescent phenotype population, and proteomic expression in oxidative stress and DNA damage repair proteins. **(A)** Quantification of BrdU cell proliferation assay after MSCs were co-treated with NAC or KU55933 (ATMi). *p < 0.05 vs. static control treatment. **(B)** Fraction of β-galactosidase-positive cells after MSCs were treated with brachial loading, NAC, or KU55933 (ATMi). *p < 0.05 vs. static control treatment. **(C)** Western blotting results and **(D)** quantification protein expression after treatment of MSCs. *p < 0.05 vs. static control treatment.

We also evaluated protein expression with western blotting pathways related to oxidative stress and DNA damage repair **(Fig. 6C-D)**. A Ponceau stain was run to confirm equal loading of cell lysate protein **(SI Fig. 2C)**. Results showed significant upregulation of SIRT6 expression with brachial and brachial + NAC treatment, but no significant change with KU55933 treatment. Furthermore, DNA damage repair protein KU80 was upregulated with brachial treatment alone but remained statistically unchanged with brachial + NAC and brachial + KU55933 treatment compared to control groups. Therefore, we concluded that key regulators of DNA damage repair such as Sirt6 and KU80 rely on ATM signaling that is activated by our mechanical treatment to protect the aged MSCs from senescence. Furthermore, mechanical enhancement of proliferative function is independent of ATM-mediated DNA damage repair signaling and instead relies on the alteration of oxidative stress signaling. Overall, these results lead to our mechanistic hypothesis that mechanical conditioning activates oxidative stress signalling through SIRT1 and antioxidant proteins such as SOD1, FOX01, and FOX03a to rejuvenate proliferative function in aged MSCs. Mechanical conditioning also activates DNA damage signalling through ATM, SIRT6, and KU80 to rapidly enhance DNA damage repair and protect the aged MSCs from the progression of senescence (**Fig 7**).

**Figure 7.**
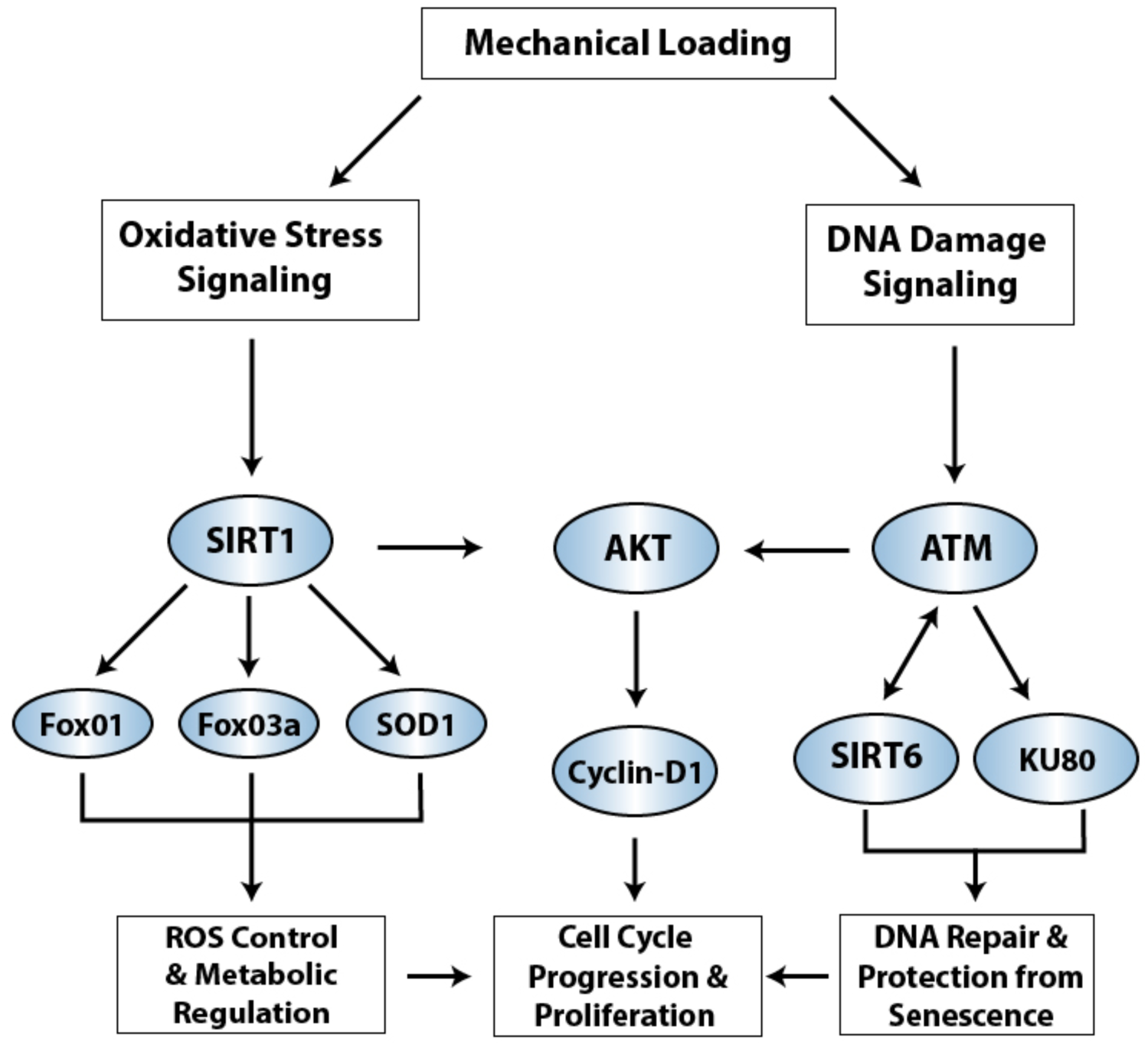
Proposed mechanistic hypothesis of how mechanical conditioning activates oxidative stress and DNA damage signaling to rejuvenate proliferative function and reduce senescence in aged MSCs.

### Mechanical conditioning reduces senescence related gene expression in MSCs from aged donors

To investigate the effects of mechanical conditioning on gene expression patterns that are correlated with senescence in MSCs, we obtained MSCs from one aged donor (81 years old) and conditioned them with brachial loading or static control treatment. Following the treatments, we performed RNA-seq analysis on the cells. A differential gene expression analysis revealed that brachial mechanical loading treatment significantly regulated 183 genes in comparison to the static control treatment **(Fig. 8A)**. Cells treated with brachial loading had similar patterns of gene expression, while MSCs treated with the static control had similar patterns of gene expression **(Fig. 8B).** Gene ontology analysis of the differential gene expression revealed significant expression increases in gene sets related to cellular metabolic processes and nucleoside biosynthetic processes **(Fig. 8C).** Previous studies have demonstrated that gene expression related to metabolic processes and protein/nucleotide biosynthetic processes are markedly downregulated in BM-MSCs from aged donors, supporting the critical role of metabolism in regulating stem cell senescence.^63^ KEGG pathway analysis further revealed a significant upregulation of biosynthesis of unsaturated fatty acids pathways in the brachial-treated group, which has been previously reported to be enriched in young, healthy MSCs **(SI Fig. 6)**.^64^

**Figure 8.**
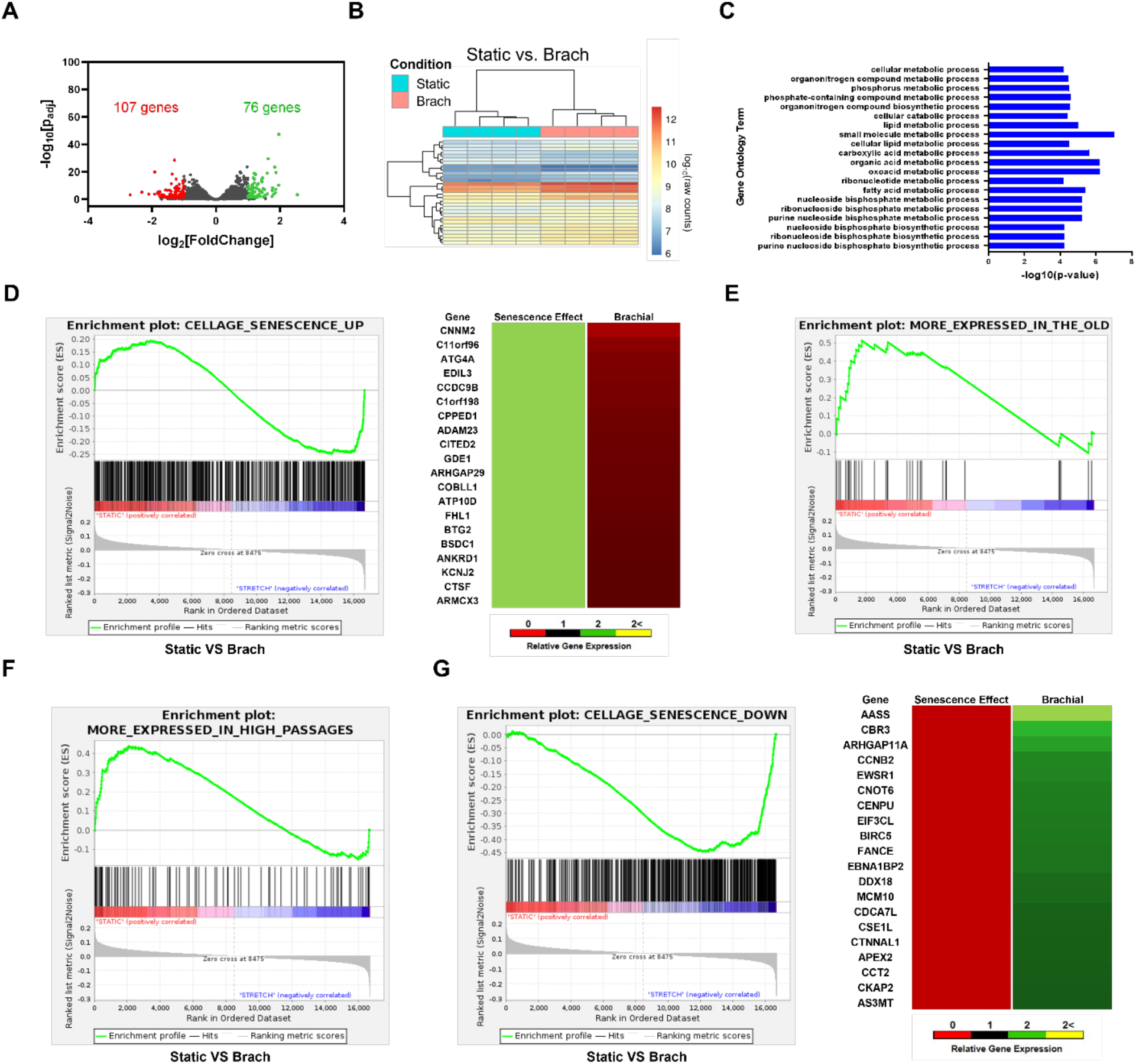
Mechanical treatment of MSCs reverses correlation of gene expression profile with senescence related gene sets. MSCs from one aged donor were conditioned with brachial mechanical loading for 7 days and analyzed with RNA sequencing. **(A)** Differential gene expression in comparison to the static control group. Genes with statistically significant upregulation (green) or downregulation (red) are shown in color. **(B)** Clustering analysis of the gene expression in the MSCs for the top significantly differentially expressed genes. *Scale:* R-log transformed values of raw gene hit counts. **(C)** Gene ontology analysis of significantly upregulated biological functional processes in the brachial-treated groups in comparison to static-treated control groups. Next, gene set enrichment analyses were performed to compare the gene enrichment in MSCs from one aged donor after control or brachial loading treatment using predefined gene sets for: **(D)** Genes upregulated in cellular senescence. FDR *q*-value = 0.013. Top 20 genes upregulated in senescence and downregulated after brachial loading. **(E)** Genes upregulated in elderly donor-derived MSCs. FDR *q*-value = 0.003. **(F)** Genes upregulated in high passage MSCs. FDR *q*-value = 0.001. **(G)** Genes downregulated in cellular senescence. FDR *q*-value = 0.0001. Top 20 genes downregulated in senescence and upregulated after brachial loading.

Next, we performed a gene set enrichment analysis (GSEA) on the RNA-seq data. We calculated the enrichment scores comparing the genes expressed between mechanically conditioned cells and control cells for gene sets upregulated in cell senescence (CellAge),^65^ upregulated in upregulated in elderly donor derived MSCs (E-MTAB-4879),^66^ or and upregulated in high passage MSCs (GSE137186).^67^ In all three senescence or aging-related gene sets, we observed a significant reduction in gene enrichment with brachial mechanical loading **(Fig. 8D-F).** We also calculated enrichment scores for a gene set downregulated in cell senescence (CellAge),^65,68^ observing a markedly significant increase in gene enrichment with brachial loading (**Fig. 8G)**. Therefore, the mechanical treatment may work to rejuvenate aged MSCs through both the suppression of age-related gene sets, as well as the enhancement of “anti-aging” gene sets, causing a broad reversal of gene expression patterns that are correlated with senescence.

To investigate the universality of this observation, we obtained MSCs from a young, healthy donor (24-year-old female) and conditioned them with brachial loading and E/E inhibitor. We repeated RNA-seq analysis on these cells and evaluated gene expression of several key gene sets related to various mechanisms of senescence in MSCs. These gene sets were identified through a thorough investigation of the literature as key mediators of MSC senescence, and the expression pattern of each gene as it pertains to MSC senescence was documented.^5,35,37,66,69–72^ The analysis found that the treatments reversed the effects implicated by senescence on genes related to age-related senescence, oncogenic-related senescence, stress-related senescence, developmental-related senescence, and inflammation-mediated senescence (**SI Fig. 7A-E**). Next, we repeated the GSEA on this RNA-seq data, and calculated enrichment scores for gene sets upregulated in high passage MSCs (GSE137186),^67^ genes upregulated in elderly donor derived MSCs (E-MTAB-4879),^66^ and genes upregulated in senescent cells (M27188).^71^ In all three senescence-related gene sets, we observed a reduction in senescence related genes with brachial mechanical loading (**SI Fig. 7F-H**).

### Mechanical conditioning modifies chromatin accessibility profile and causes enrichment of transcription factor motif binding in aged MSCs

To investigate the mechanical loading effect on altered chromatin accessibility, we obtained MSCs from one aged donor (81 years old) and conditioned them with brachial loading or static control treatment. Following the treatments, we performed an assay for transposase-accessible chromatin using sequencing (ATAC-seq) on the cells. We compared the enrichment of accessible chromatin at genomic features between static and brachial treated cells **(Fig. 9A)**. A differential analysis of accessible chromatin peaks revealed 50 genomic regions that were significantly more accessible in static treated cells, and 293 genomic regions significantly more accessible in brachial treated cells. The top 20 most differentially upregulated and downregulated ATAC-seq peaks for brachial vs. static treated cells are listed in **Fig. 9B**. A Reactome pathway enrichment analysis of biological functions most associated with highly accessible chromatin regions revealed upregulated MAPK family signaling and antigen processing functions with brachial loading treatment **(Fig. 9C)**. Next, we used the Hypergeometric Optimization of Motif EnRichment (HOMER) tool to identify transcription factor motif enrichment of accessible chromatin peaks in brachial treated cells **(Fig. 9D-F)**.^73^ We classified the enriched transcription factor motifs based on their biological function and interaction with other regulatory factors. We identified several related transcriptional regulators involving the AP-1 complex **(Fig. 9D)**.^74^ Factors from this list, including c-Jun and AP-1 have been shown to act as co-activators of DNA methyltransferase 1 (DNMT1) to inhibit senescence progression and increase differentiation potential in MSCs.^75^ We also identified transcription factor motifs involved in mechanosignaling pathways, including members of the SMAD and TEAD family, whose enrichment was upregulated with brachial loading **(Fig. 9E).** Notably, several of these factors were identified in our previous study as potential drivers of mechanical conditioning-induced signaling to improve MSC proliferation and vascular regeneration.^31^ We also observed several transcription factor motif families, including RUNX, PAX, and GATA, that are known to regulate resident stem cell development, proliferation, and differentiation potential in skeletal muscle and cardiac tissue **(Fig. 9F)**.^76^ Lastly, we integrated the ATAC-seq results with the RNA-seq data to identify potential relationships between chromatin accessibility and differential gene expression **(Fig. 9G, H)**. We mapped transcription start sites to the top 20 differentially enriched peaks for **(G)** brachial treated cells and **(H)** static control treated cells, then compared the differential expression of these genes for brachial vs. static treated groups. Upregulated genes include ALCAM, a known marker of normal MSC phenotype, and GPAM, a regulator of healthy mitochondrial metabolism in MSCs.^77^ Downregulated genes include BRD4, a transcriptional co-activator that activates oncogene-induced senescence and SASP signaling.^78^

**Figure 9.**
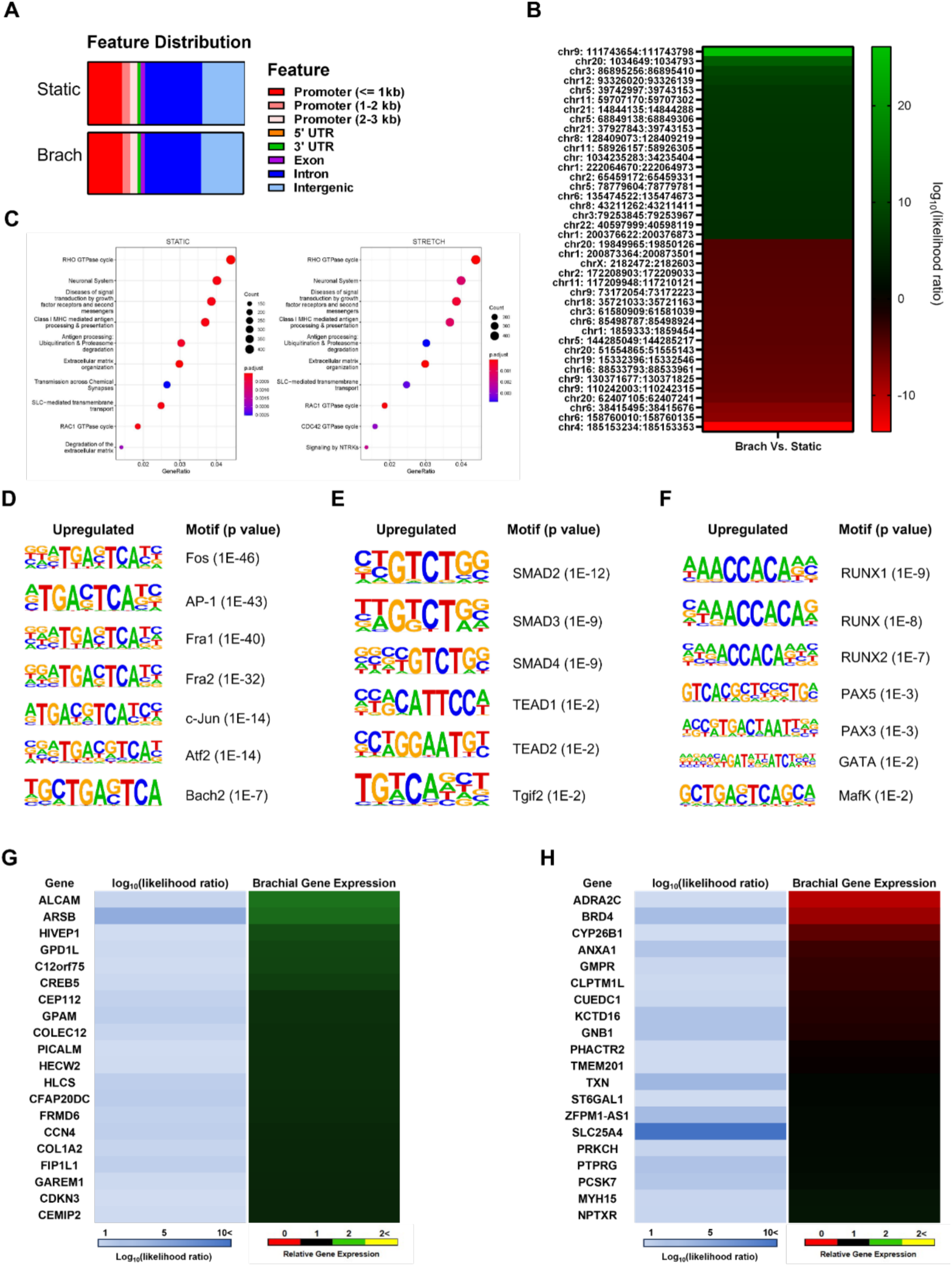
Mechanical conditioning of aged MSCs causes altered chromatin accessibility profiles. MSCs from one aged donor were conditioned with brachial mechanical loading for 7 days and analyzed with ATAC sequencing. **(A)** Enrichment of accessibility chromatin peaks at genomic features. **(B)** Heatmap of top 20 most differentially upregulated and downregulated ATAC-seq peaks at accessible chromatin regions in brachial vs control treated cells. **(C)** Reactome pathways associated with highly accessible chromatin regions. **(D-F)** HOMER transcription factor motif enrichment analysis of the upregulated ATAC-seq peaks in brachial treated cells. **(D)** Transcriptional regulators involving the AP-1 complex. **(E)** Mechanosensitive signaling factors involving the TGF-β –SMAD2/3 and Yap/Taz-TEAD pathways. **(F)** Transcriptional regulators of MSC phenotype and resident stem cell proliferation and differentiation potential in skeletal muscle and cardiac tissue. **(G, H)** Integrated ATAC-seq and RNA-seq analysis. Transcription start sites were mapped to the top 20 differentially enriched peaks for **(G)** brachial treated cells and **(H)** control static treated cells. The differential gene expression of these genes for brachial treated samples was compared with differential chromatin accessibility.

## DISCUSSION

Mesenchymal stem cells are a promising cell type for autologous regenerative therapies, but clinical trials have largely yielded inconsistent and/or disappointing results. A major potential reason for these limitations is cellular senescence, caused by poor donor health, advanced age, and extensive *in vitro* expansion required to generate an adequate number of cells for therapeutic applications.^5^ In our study, we mechanically conditioned MSCs derived from aged donors to enhance their short and long-term proliferative function, improve their multipotency, and protect the cells from advancing senescence, resulting in a lower population of senescent cell phenotypes following the treatment. Our study extends previous work conducted by our laboratory, in which we demonstrated that brachial mechanical loading in co-treatment with an EGFR/ErbB-2/4 (E/E) synergistically enhanced the angiogenic properties of MSCs.^28^ Our findings in this study support that brachial loading alone was superior in enhancing aged MSC proliferative function, expansion capacity, and multipotent properties. For application in vascular regenerative medicine, a combination of the two approaches could provide a powerful approach to improving the effectiveness of MSCs. Brachial loading alone could be used to rejuvenate the cells and allow their expansion while brachial loading with the E/E inhibitor would provide a step to induce vascular phenotype, production of soluble HGD and enhance their therapeutic efficacy for treating ischemia and inducing revascularization. The brachial loading as a first step to MSC expansion could also be useful in many other application if later followed by differentiation into other phenotypes.

One interesting question that arises from our study is the potential universality of the mechanical loading regime to improve cell culture expansion of many other cell types. Our computational model would suggest that mechanical strains in the correct range lead to forces on the chromatin that could lead to opening of the DNA and that the brachial loading, in particular, has an essential a wider range where the DNA could be opened without being damaged.^50^ Combined with our studies here, this would suggest that many cell types would benefit from similar biophysical treatment to enhance their culture expansion, reduce senescence and induce DNA damage repair. To investigate this concept, we treated high-passage (P10) human umbilical vein endothelial cells with brachial loading and performed assays to test their proliferation, β-Galactosidase expression, and long-term culture expansion capacity (**SI Fig. 8**). Results demonstrated that brachial conditioning did not improve the cells’ short-term DNA synthesis but did significantly reduce the population of β-Galactosidase+ cells and improved their cumulative doubling rate over long-term *in vitro* expansion. Future studies may consider the generation of cell-type-specific mechanical conditioning regimes to improve therapeutic efficacy for a wide range of disease conditions and regenerative applications.

Other studies have shown that the application of mechanical forces can induce antioxidant response and improve the proliferative potential of MSCs.^79,80^ Thus, there is rationale that specific types of mechanical conditioning may have a rejuvenating effect on MSCs, although the mechanism remains largely unclear. Our work demonstrated that brachial mechanical loading resulted in the upregulation of several Sirtuin proteins, which are known to be crucial regulators of aging. Several current clinical trials are investigating the anti-aging potential of pharmacological SIRT1 activators such as Resveratrol^81^ and Metformin,^82^ although results remain inconclusive in human subjects. Our work demonstrates a potential mechanism by which Sirtuin proteins alter key pathways of senescence, oxidative stress management and DNA damage repair, to rejuvenate the functionality of aged MSCs. Previous work has shown that SIRT6 functions as a DNA damage sensor and activates DNA damage signaling through the recruitment and phosphorylation of ATM and γ-H2AX histone foci.^83^ The activated ATM in turn phosphorylates Akt to promote cell survival and proliferation,^84^ which is in agreement with the results of our mechanistic studies **(Fig. 7)**.

We investigated the novel finding that mechanical conditioning of aged MSCs results in the enhanced recognition and repair of DNA double-strand-breaks. We found that enhanced DNA damage repair is not dependent on Piezo1-mediated mechanosensitive signaling, although the inhibition of Piezo1-related pathways did prevent a subset of double strand break repair. Next, we investigated oxidative stress and DNA damage repair pathways for proteomic signaling, finding that mechanical conditioning resulted in the broad activation of antioxidant proteins such as SOD1, FOX01, FOX03a and FOX04, and DNA damage repair proteins such as KU80 and ATM. We also evaluated the downstream functional effects of the alteration of these pathways, finding that mechanical conditioning reduced the buildup of reactive oxygen species and resulted in rapid repair of DNA damage. When we altered oxidative signaling through the co-treatment of aged MSCs with the antioxidant N-Acetyl Cysteine (NAC) and brachial loading, we observed a reduction of proliferative enhancement. Conversely, when we altered DNA damage repair signaling through the co-treatment of ATM inhibitor KU55933 with brachial loading, we observed a reduction of senescence-protective effects, resulting in a larger phenotype of β-Galactosidase+ senescent cell phenotypes. Together, these results suggest that mechanical conditioning relies upon altered signaling in both key pathways of senescence to rejuvenate specific aspects of the cells’ functionality.

Several studies have targeted the elevated reactive oxygen species (ROS) and impaired antioxidant mechanisms that are characteristic of senescent MSCs, using gene vectors to overexpress the sirtuin proteins,^26,85^ antioxidant pharmaceutical treatment,^86,87^ or conditioning with extracellular exosomes.^27^ These methods have provided enhancement in MSC resistance to apoptosis and oxidative stress, improvement in functionality in in vivo models, and increases in MSC growth. Our work has examined using stretch as a biophysical stimulus for reducing senescence and DNA damage. In our studies, the biophysical stimulation activated antioxidant pathways and sirtuins, in addition to many other pathways, similar to those used to reduce senescence in MSCs in prior studies. A key benefit of our approach was the activation of multiple pathways with the biophysical stimulus and the potential direct benefit of inducing DNA damage repair in chromatin through mechanical stretching, in the absence of pharmaceutical treatment or genetic modification. Perhaps the most striking aspect of our findings is the long-term effects of a relatively short mechanical protocol. One week of loading at 4 hours per day lead to long-term changes in expansion potential at least 8 passages later and reduction in senescence. In this work, we only explored one week of conditioning and further work could explore whether multiple rounds of this treatment could further enhance expansion.

Overall, our study demonstrated that mechanical conditioning led to upregulated antioxidant expression and enhanced DNA damage repair, suggesting that there is a crucial homeostatic balance of signaling in and between these pathways in the rejuvenation of senescence. Therefore, while highly targeted attempts to improve the antioxidant or DNA damage repair response genetically or pharmacologically may cause unintended consequences, our mechanical treatment resulted in the broad enhancement of these pathways in aged MSCs to restore their functionality. Furthermore, by investigating the enrichment of key gene sets related to senescence, our study demonstrated that mechanical conditioning alters chromatin accessibility and transcriptional machinery to reverse the expression patterns of age-related gene sets, while enhancing the expression patterns of gene sets known to be downregulated in senescence. These key findings illustrate the potential for generating long-term, lasting benefits in MSC functionality following a week-long mechanical conditioning period. We propose that our biomechanical conditioning may generate an restored chromatin state to suppress cryptic and dysregulated transcription patterns common in aged stem cells.^88^ Taken together, these findings may provide a practical method to rejuvenate the essential homeostatic functions of MSCs from aged patients to improve their expansion for use in therapeutic applications.

## Acknowledgements.

The authors gratefully acknowledge funding through the DOD CDMRP (W81XWH-16-1-0580; W81XWH-16-1-0582) and the National Institutes of Health (1R01HL141761-01) to ABB.

## MATERIALS AND METHODS

### Cell lines and cell culture

Human mesenchymal stem cells (Promocell) were cultured in low glucose DMEM medium supplemented with 15% fetal bovine serum, L-glutamine and penicillin/streptomycin. Following trypsinization, cells were seeded on the membranes at 20,000 cells per cm^2^ before mechanical loading. The cells were used between passages 4-6, unless otherwise specified. The MSCs were derived from a donor 1 (Caucasian female, 81 years old), unless otherwise specified. A subset of the studies were performed using MSCs from donor 2 (Caucasian male, 63 years old), or donor 3 (Caucasian male, 68 years old), or donor 4 (Caucasian female, 92 years old). A subset of the studies used MSCs derived from a young, healthy donor (Caucasian female, 24 years old). Human umbilical vein endothelial cells (HUVECs; PromoCell GmbH) were cultured in MCDB 131 medium supplemented with 7.5% fetal bovine serum, L-glutamine and penicillin/streptomycin and EGM-2 SingleQuot Kit (Thermo Fisher Scientific, Inc.). All cells were cultured in an incubator at 37ᵒC under a 5% CO_2_ atmosphere.

### Application of biomechanical treatment

Mechanical strain was applied to cell culture using a high throughput system described previously.^32,89,90^ Briefly, cells are cultured on custom made plates that are mounted on a system that applies strain. The cell culture plates were comprised of silicone membranes (0.005” thickness; Specialty Manufacturing, Inc.) that are sandwiched between two plates, with silicone rubber gaskets at the interfaces to prevent leaking. These cell culture membranes are UV sterilized and coated with 50 µg/mL fibronectin overnight at 37°C to allow cell adhesion. After cell attachment, the plates are mounted on to the top plate of the system using screws. To apply mechanical strain, a platen with 36 Teflon pistons is moved into the cell culture membrane. The motion is driven by a hygienically sealed, voice coil-type linear motor (Copley Controls). The platen is stabilized using six motion rails mounted with linear motion bearings. The hygienically sealed motor housing has chilled water running through in order to prevent overheating during operation. Unless otherwise specified, loading was applied for 4 hours per day for 7 days with or without 1 µm the EGFR/ErbB-2/ErbB-4 Inhibitor (CAS 881001-19-0; **Supplemental Table 3**).

### BrdU Proliferation Assay

Following the mechanical and chemical treatments, cells were removed from the stretch plates using 0.05% Trypsin-EDTA, seeded onto optical-bottom 96 well plates at a density of 10,000 cells/well, and allowed to grow for 24 hours. Bromodeoxyuridine (5-bromo-2’-deoxyuridine; BrdU) labeling solution was added to culture media. Cells were washed and stained with the detection and HRP antibodies, following the BrdU assay protocol according to the manufacturer’s instructions (Cell Signaling Technology). Absorbance was read at 450 nm using a FlexStation-3 plate reader (Molecular Devices).

### Long-term culture expansion assay

Following the mechanical and chemical treatments, cells were removed from the stretch plates using 0.05% Trypsin-EDTA and counted with a hemacytometer. Cells were seeded into standard 6 well culture plates (Corning) at a density of 100,000 cells/well, with an n=2 for each treatment group. Every 7 days, cells were trypsinized, counted, and reseeded into new 6 well plates at an equal cell seeding density. At the end of the study, cumulative doubling events across all passages were calculated for each treatment group.

### Cell lysis and immunoblotting

Following the treatments, the cells were lysed in 20 mM Tris with 150 mM NaCl, 1% Triton X-100, 0.1% SDS, 2 mM sodium orthovandate, 2 mM PMSF, 50 mM NaF and a protease inhibitor cocktail (Roche, Inc.). The proteins were separated on a NuPAGE 10% bis-tris midi gel (Novex) and transferred to nitrocellulose membrane using iBlot transfer stack (Novex). The membranes were blocked for one hour in 5% non-fat milk in PBS with 0.01% tween-20 (PBST). After washing twice in PBST, cells were incubated with primary antibodies (**Supplemental Table 1**) overnight in 1% non-fat milk at 4°C. The membranes were washed with PBST and incubated at room temperature for two hours with secondary antibody. The membrane was treated with chemiluminescent substrate (SuperSignal West Femto; Thermo Fisher Scientific, Inc.) then imaged using a digital imaging system (Cell Biosciences, Inc.).

### Human mesenchymal stem cell multipotency analysis

MSCs at passage 4 were cultured and treated with biomechanical conditioning as described previously. Following the treatments, MSCs removed from the stretch plates with 0.05% Trypsin-EDTA and seeded onto standard culture plates, then expanded under normal culture conditions until passage 8. Following the protocol from the hMSC functional identification kit (R&D Systems; #SC006), P8 MSCs were removed from the culture plates with 0.05% Trypsin-EDTA and seeded onto glass coverslips in 24 well plates at a density of 2.1×10^4^ cells/cm^2^ for the adipogenic differentiation and 4.2×10^3^ cells/cm^2^ for the osteogenic differentiation. Cells were treated with the respectively provided differentiation media every 3 days for a total of 21 days. Following the differentiation, the cells were fixed in 4% paraformaldehyde in PBS for 10 minutes followed by washing with PBS. Next, samples were blocked and permeabilized with PBS containing 5% FBS, 1% BSA, and 0.3% Triton X-100 PBS for 40 minutes. After washing, cells were incubated with primary antibodies (see **Supplemental Table 2** for specific antibodies and concentrations) in PBS with 1% BSA overnight at 4ᵒC. The samples were then washed twice in PBS with 1% BSA and incubated with secondary antibodies in PBS with 1% BSA for 2 hours in a light protected environment. Cells treated with extensive washes with PBS with 1% BSA prior to flipping the coverslip and mounting on glass slides in anti-fade media (Vector Laboratories, Inc.). The samples were then imaged using confocal fluorescence microscopy (LSM 710 Confocal; Carl Zeiss, Inc.).

### Reactive oxygen species assay

MSCs were treated with brachial mechanical loading for 2 hours. Next, culture media was removed, and chemical treatments were added to fresh cell media. Cells received hydrogen peroxide (H2O2; 0.4 mM in PBS), N-acetyl-cysteine (NAC; 5 mM in culture media), or fresh culture media only. The mechanical loading was resumed for 1 additional hour, and chemical treatments were replaced with fresh culture media. The mechanical loading was then resumed for an additional 30 minutes. 4 µL of CellRox DeepRed reagent (ThermoFisher Scientific) was added directly to the cell culture media in each well, and the mechanical loading was resumed for an additional 30 minutes. Following the loading, cells were washed with PBS, removed from the stretch plates using 0.05% Trypsin-EDTA, and centrifuged for 3 minutes at 300G, and supernatant removed. Cells from each treatment group were resuspended in 1 mL room temperature stain buffer (BD Biosciences) in flow cytometry tubes (Falcon). Cell suspensions were centrifuged for 3 minutes at 300G, supernatant removed, and resuspended in 1mL of stain buffer with 1 µL of Sytox Blue Live/Dead reagent (ThermoFisher Scientific) and incubated for 10 minutes at room temperature in a light-protected environment. Cell suspensions were centrifuged, supernatant removed, and resuspended in 1mL of 1X fix/perm solution (BD Biosciences) and incubated at 4ᵒC for 30 minutes. Following fixation, cells were centrifuged with washing buffer two more times, before resuspended in stain buffer and measured. A BD LSR II Fortessa Flow Cytometer (BD Biosciences) was used to measure population fluorescent signals. At least 10,000 events were recorded and further gating and quantification was done through FlowJo software.

### Immunocytochemical staining on silicone membranes

Following the treatments, the cells were fixed in 4% paraformaldehyde in PBS for 10 minutes followed by washing and permeabilization with 0.1% Triton X-100 PBS for 5 minutes. Next, samples were blocked with PBS containing 5% FBS and 1% BSA for 40 minutes. After washing, cells were incubated with primary antibodies (see **Supplemental Table 2** for specific antibodies and concentrations) in PBS with 1% BSA overnight at 4ᵒC. The samples were then washed twice in PBS with 1% BSA and incubated with secondary antibodies in PBS with 1% BSA for 2 hours in a light protected environment. Cells treated with extensive washes with PBS with 1% BSA prior to mounting in anti-fade media (Vector Laboratories, Inc.). The samples were then imaged using confocal microscopy with a LSM 710 laser scanning confocal microscope (Carl Zeiss, Inc.).

### p-γ-H2AX Foci Analysis

Following confocal imaging, images were analyzed with FIJI software to count p-γ-H2AX histone foci. Three-dimensional image stacks were maximum intensity projected to generate two-dimensional images with 12-bit range of 0-4095. Background signal outside of nuclei boundaries was measured and subtracted from the image. Individual cell nuclei regions of interest were selected, and an intensity threshold of 300 bits was applied. The Analyze Particle plugin was applied to count the p-γ-H2AX foci in each nucleus, with a minimum particle size of 0.5 µm applied. At least 100 individual cell nuclei were analyzed for each treatment group to calculate the mean foci count per cell, mean foci count per nuclear area, and standard errors.

### Nuclear Morphometric Analysis

Following confocal imaging, DAPI images were analyzed with the Nuclear Irregularity Index (NII) plugin in FIJI to analyze the nuclear morphology of the cells.^51^ Nuclei were manually marked by defining the nuclei intensity segmentation threshold, as described in the NII plugin manual. Images of nuclei from a young, healthy (24-year-old) MSC line were used to parametrize the “normal nuclei”. Images of nuclei from brachial or static control treated MSCs were included as “treated nuclei”, with at least 100 nuclei analyzed for each group. Nuclei were classified as “Normal”, “Apoptotic”, “Senescent”, or “Irregular” based on measurements of their area, area/box, aspect, radius ratio, and roundness, in comparison to the young “normal nuclei” control.

### Comet assay

Cells were treated with brachial mechanical loading at 0.1 Hz and 7.5% maximum strain for 30 minutes, 4 hours, cultured under static conditions, or cultured under static conditions with 0.4 mM H2O2 to induce DNA damage. Following the treatments, a subset of the cells were left in static conditions for 20 hours. Next, a neutral comet assay was performed to quantify the DNA double-strand-breaks, using the Comet Assay Kit (Abcam), and following the manufacturer’s protocol. Briefly, the lysis buffer, alkaline solution, and TBE neutral electrophoresis solution were prepared in advance and stored at 4ᵒC. Comet agarose solution was heated at 90ᵒC for 20 minutes, stirred until homogenized, and pipetted onto the comet slides, then stored horizontally at 4ᵒC. Following the treatments, culture media was removed from the cells, then cells were washed with dPBS and incubated with 25 mM EDTA for 10 minutes at 37ᵒC. Cells were gently removed with pipetting, centrifuged, and resuspended at 2×10^5^ cells/mL in 4ᵒC dPBS. Cell suspensions were combined with molten comet agarose at a 1/10 (v/v) ratio. 75 µL of cell/agarose suspensions was pipetted into each well of the comet slides, on top of the agarose base layer. Slides were allowed to chill at 4ᵒC until the agarose solidified. Next, slides were incubated with 4ᵒC lysis buffer for 60 minutes in light-protected conditions. Lysis buffer was aspirated and replaced with 4ᵒC alkaline solution and incubated for 30 minutes in light-protected conditions. Alkaline solution was aspirated and replaced with 4ᵒC TBE electrophoresis solution, then slides were carefully transferred to a horizontal electrophoresis chamber (BioRad). 20 volts were applied for 20 minutes, then slides were transferred to 4ᵒC deionized water for washing. Lastly, slides were fixed with 4ᵒC 70% ethanol, then air dried. 1X Vista Green DNA Dye (Abcam) was added to the slides and incubated at room temperature for 15 minutes. The samples were then imaged using confocal microscopy with a LSM 710 laser scanning confocal microscope (Carl Zeiss, Inc.). The OpenComet plugin in FIJI was used to score the comets and quantify the Olive Tail Moment (OTM) for each treatment group,^91^ with at least 100 comets scored per group.

### Conditioning of hMSCs using biochemical factors

For mechanistic experiments involving chemical factors and biological inhibitors, cells were incubated with the chemical treatments as shown in **Supplemental Table 3.** The cells were treated with mechanical loading for 4 hours/day under brachial waveform at 0.1 Hz and 7.5% maximum strain or cultured under static conditions. The culture media containing the treatments were replaced on day 3 and day 5 for all 7-day treatments.

### Β-Galactosidase Analysis

Following the treatments, cell culture media was removed, and cells were washed once with PBS. Cells were fixed and stained for β-Galactosidase following the protocol of the Senescence β-Galactosidase Staining Kit (Cell Signaling Technologies). Following staining, cells were stored with 70% glycerol and imaged using a light microscope (Meiji, Inc.). Images were post-processed in Adobe Photoshop to increase image contrast and visibility of β-Galactosidase+ cells. Total cells and β-Galactosidase+ cells were manually counted in FIJI, with at least 1,000 cells counted per treatment group.

### Flow Cytometry

For the characterization of mesenchymal stem cells during long-term culture expansion (passage 5-10), the cells were detached from the culture plate using Accutase (Sigma-Aldrich) and were labeled with fluorescent antibodies according to the R&D Systems human mesenchymal stem cell verification flow kit protocol (R&D FMC020; see **Supplemental Table 4** for the specific antibodies used). Briefly, the detached cells were centrifuged and the supernatant was removed. Fixing and permeabilizing buffer was added while the cells were vortexed and incubated for 40 min. Next, the samples were centrifuged and the supernatant was removed. Cells were then treated with washing buffer containing antibodies for 50 min. Following antibody incubation, cells were centrifuged with washing buffer two more times, then were treated with stain buffer and measured. A BD LSR II Fortessa Flow Cytometer (BD Biosciences) was used to measure population fluorescent signals. At least 10,000 events were recorded and further gating and quantification was done through FlowJo software.

### RNA Sequencing and Analysis

Following treatments, RNA was isolated from the cells from four independent wells per group using the Qiagen RNeasy Mini Kit. RNAseq was performed using an Illumina HiSeq 2500 sequencing machine. For sequencing, single reads of 50 base pairs were performed after poly-A mRNA capture using the Poly(A) Tailing Kit (Ambion) and Ultra II Directional RNA Library Prep Kit (NEB) to isolate mRNA and perform dUTP directional preparation of the mRNA library. RNA sequencing was performed by GENEWIZ (Azenta Life Sciences). Gene expression analysis was performed using R. Gene ontology, Pathview, and Kyoto Encyclopedia of Genes and Genomes (KEGG) analysis was performed using the Molecular Signatures Database (En), PANTHER, or ShinyGO.^92–94^

### Gene set enrichment analysis (GSEA)

To define signature gene sets of senescent cells, aged-donor-derived MSCs, or high passage MSCs, we downloaded multiple RNA-seq data sets **(Supplemental Table 5).** Obtained RNA-seq data sets were mapped to the human transcripts (GRCh38) with Salmon mapper (v1.1.0)^95^ and the count of mapped reads was normalized as transcripts per million (TPM) using an R package of tximport (v 1.14.2)^96^. Multiple processed RNA-seq data belonging to each cell-type category were considered as replicates and compared to data obtained from mesenchymal stem cells (MSCs) using edgeR (v 3.26.8)^97^ to define signature genes for each cell-type. GSEA was performed using fig 4 tool (v4.0.3)^98^ with these defined gene sets (p-value 0.001, approximately 500 genes each) and the gene expression data of the brachial treated cells to control static treated cells.

### Statistical analysis

All results are shown as mean ± standard error of the mean. All experiments used biological replicates that consisted of cells in non-repeated, independent cell culture wells or tissue samples from different animals, unless specified otherwise. Multiple comparisons between groups were analyzed by two-way ANOVA followed by a Tukey post-hoc or a Dunnett post-hoc test when testing multiple comparisons versus a control group. For non-parametric data, multiple comparisons were made using the Kruskal-Wallis test followed by post-hoc testing with the Conover-Iman procedure. A *p*-value of 0.05 or less was considered statistically significant for all tests.

## SUPPLEMENTAL FIGURES

**Supplemental Figure 1.**
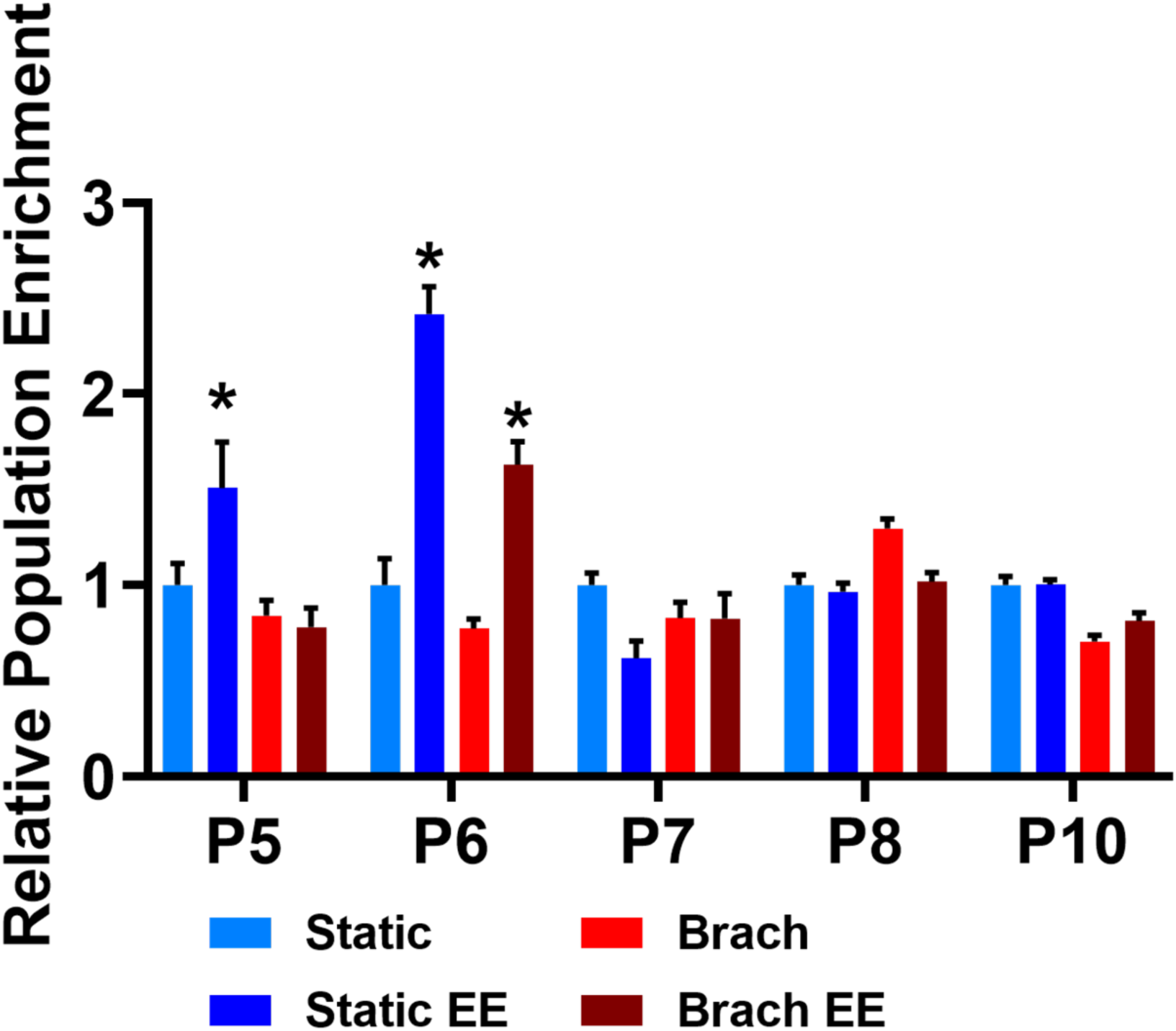
Verification of MSC phenotype in MSCs after biomechanical conditioning during passage 5 and subsequent culture expansion. The negative marker cocktail included antibodies for CD45, CD34, CD11b, CD79A and HLA-DR.

**Supplemental Figure 2.**
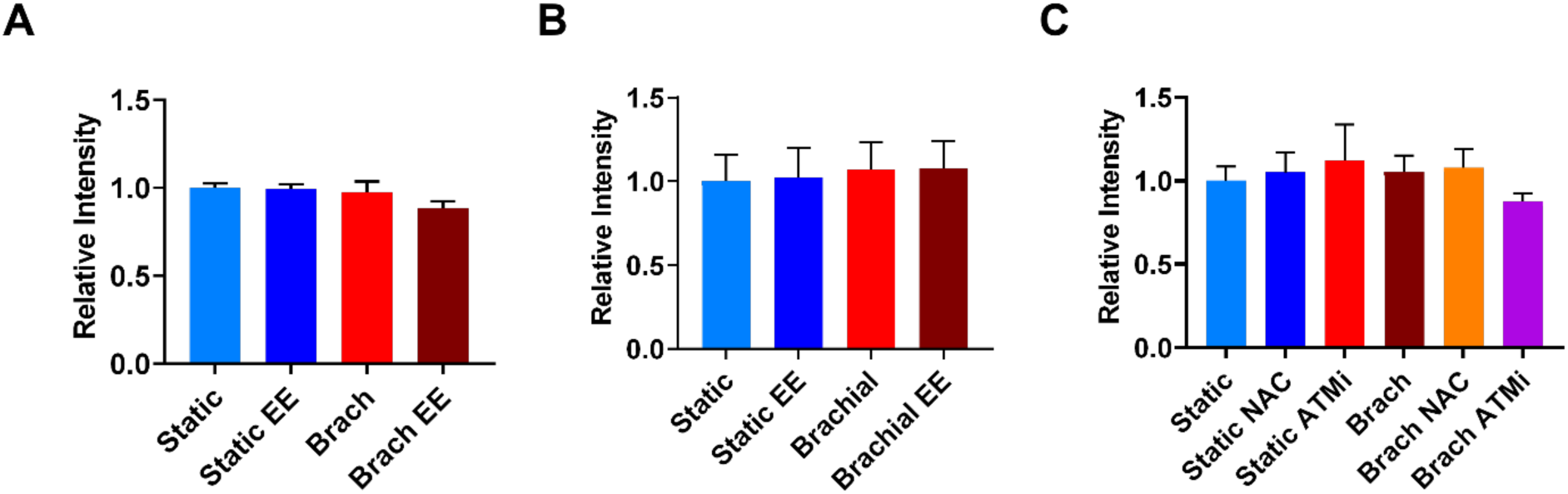
Ponceau staining quantification to confirm equal loading of cell lysate protein.

**Supplemental Figure 3.**
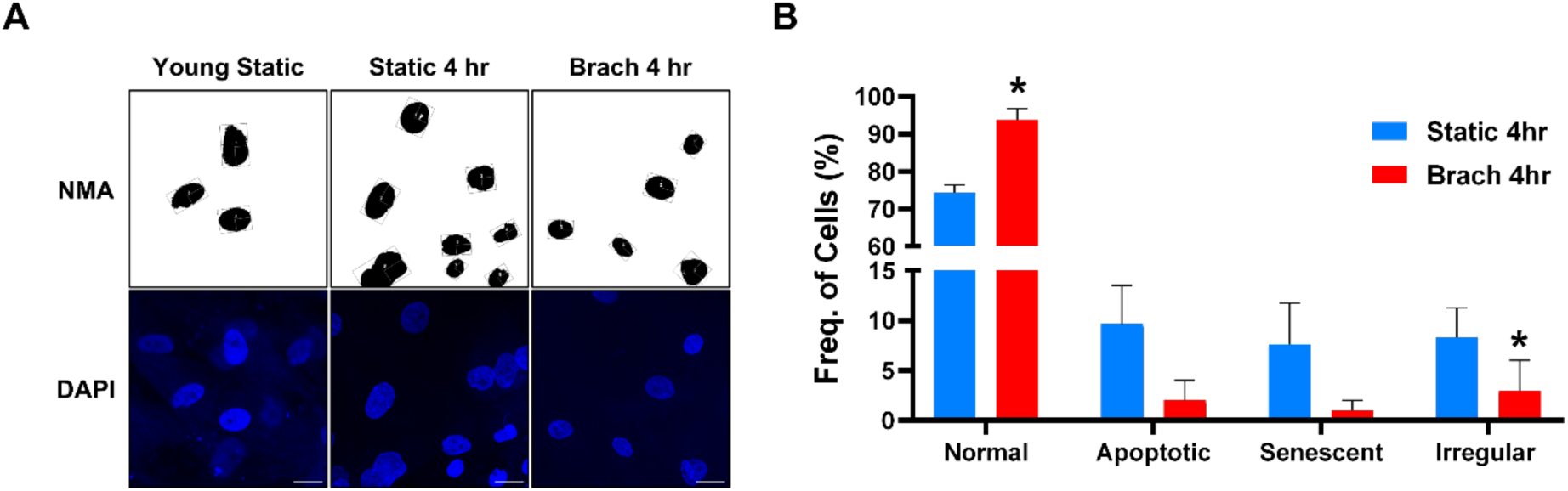
MSCs from a young donor (24 years old) or aged donor (81 years old) were treated with brachial loading or static control treatment for 4 hours and stained for DAPI. The Nuclear Irregularity Index (NII) plugin in FIJI was used to assess the nuclear morphology of the cells. **(A)** Sample images processed with the NII plugin. Scale bar = 20 µm. **(B)** Quantification of nuclear morphometric analysis. Nuclei were scored as “Normal”, “Apopotic”, “Senescent”, or “Irregular”. *p < 0.05 vs. static control treatment.

**Supplemental Figure 4.**
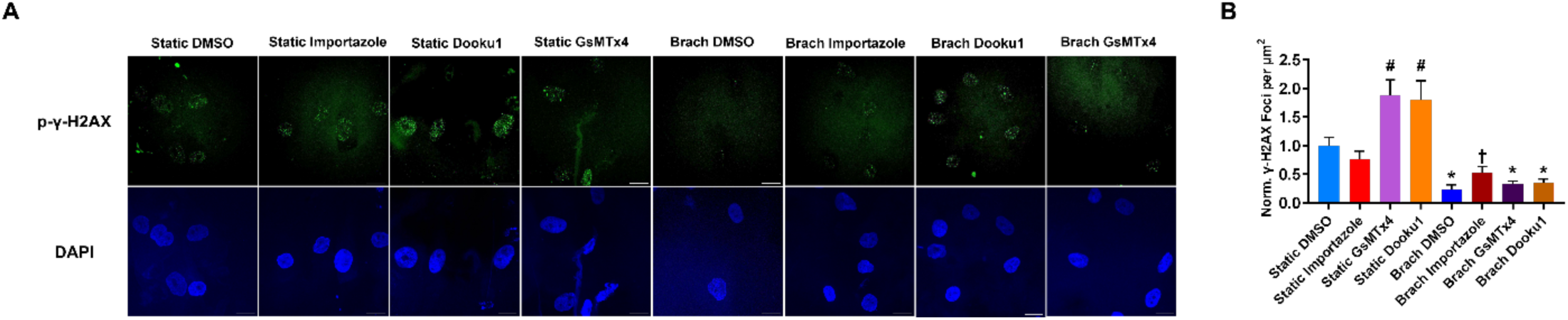
MSCs were treated with brachial loading or static control treatments for 4 hours and co-treated with 40 µM Importazole, 2 µM Dooku 1, 3 µM GsMTx4, or DMSO. **(A)** Immunostaining of p-γ-H2AX histone foci and DAPI. Scale bar = 20 µm. **(B)** Quantification of p-γ-H2AX histone foci expression per nuclear area. *p < 0.05 vs. matching static treatment. #p < 0.05 vs. static control treatment. †p < 0.05 vs. brachial DMSO treatment.

**Supplemental Figure 5.**
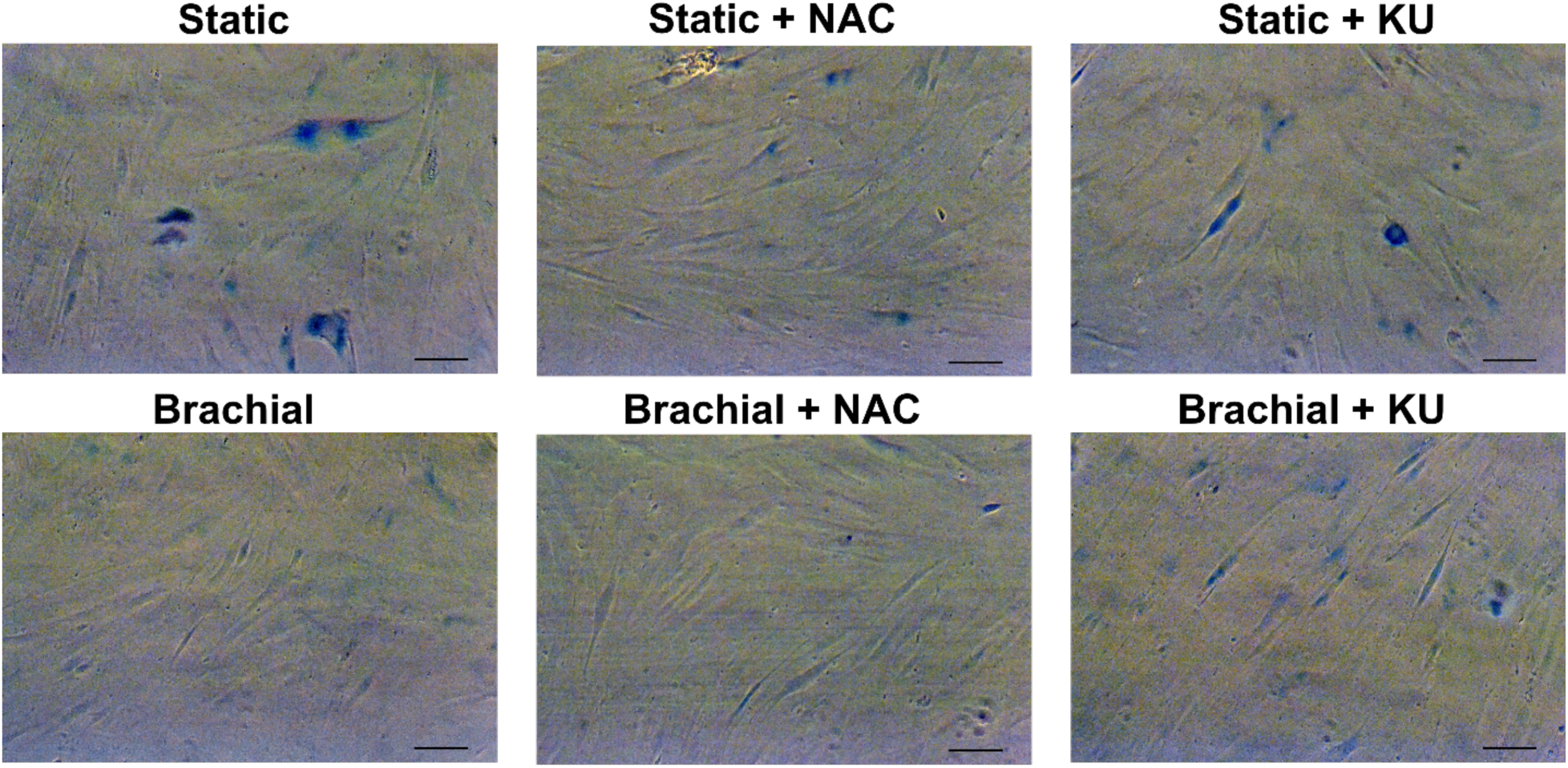
Images of β-galactosidase staining after MSCs were treated with brachial loading, NAC, or KU55933 (ATMi). Scale bar = 100 µm.

**Supplemental Figure 6.**
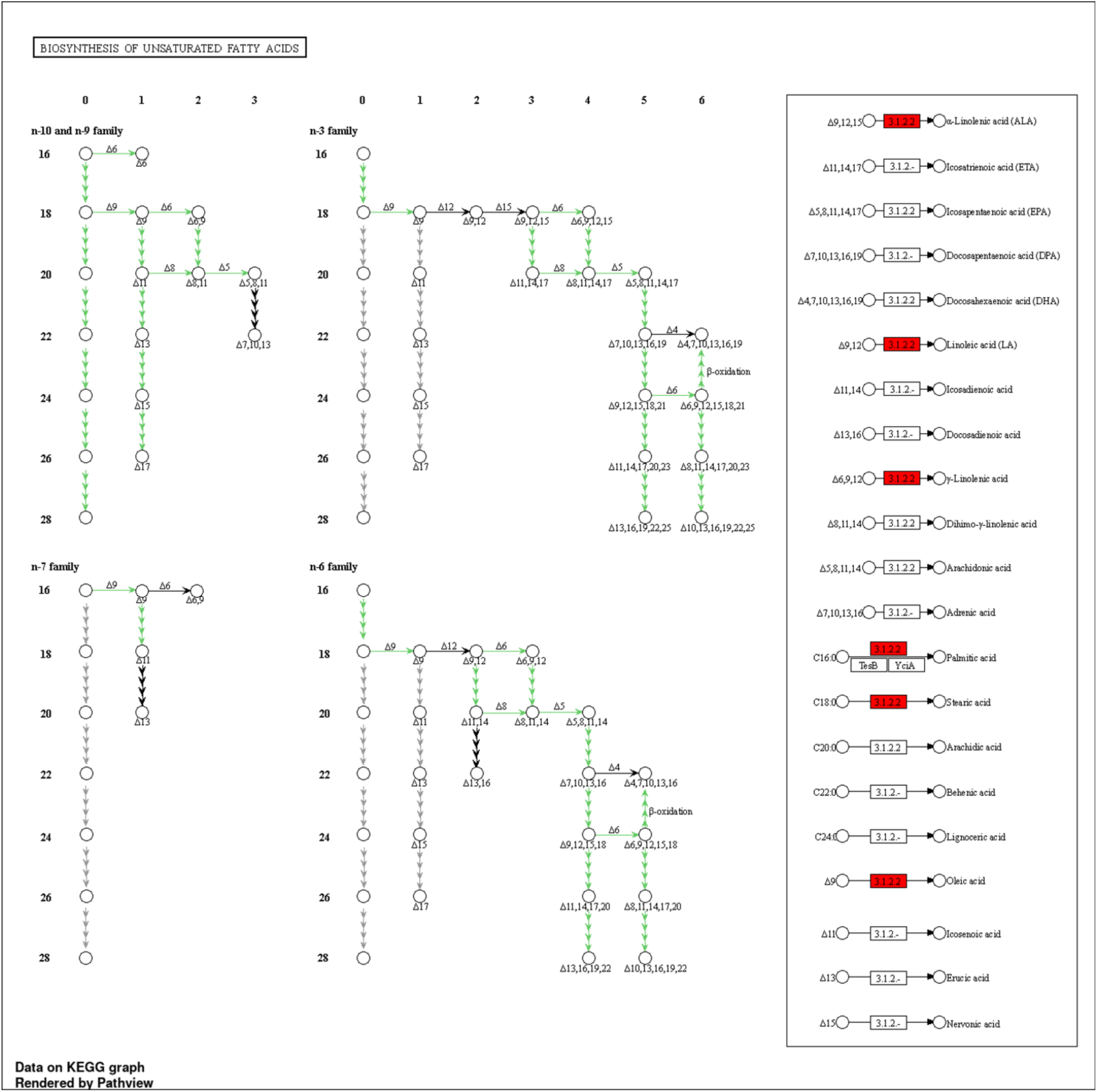
KEGG pathway analysis demonstrating significant upregulation of biosynthesis of unsaturated fatty acids-related pathways in brachial treated MSCs.

**Supplemental Figure 7.**
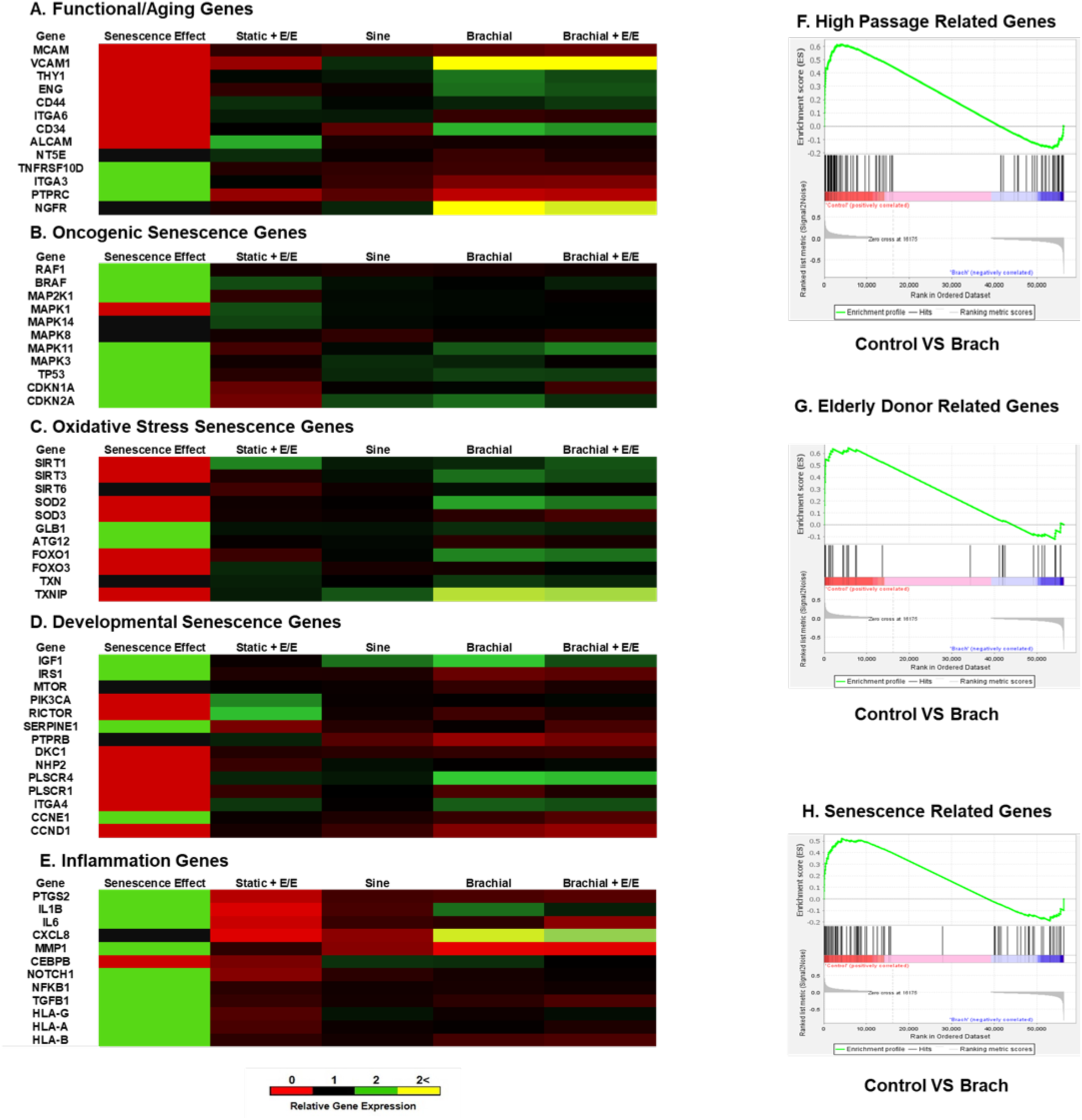
MSCs from a young donor (24 years old) were conditioned with mechanical and pharmaceutical treatments and analyzed with RNA sequencing.^31^ A transcriptomic analysis was performed to compare the gene expression of treated MSCs with the gene expression patterns of key senescence-related genes in MSCs. Results revealed the reversal of senescence-related gene expression patterns with mechanical and pharmaceutical treatment in gene sets for: **(A)** Aging senescence-related genes. **(B)** Oncogenic senescence-related genes. **(C)** Oxidative stress senescence-related genes. **(D)** Developmental senescence-related genes. **(E)** Inflammation-mediating genes. Next, a Gene Set Enrichment Analyses was performed to compare the gene enrichment in MSCs from young donors after control or brachial loading treatment using predefined gene sets for: **(F)** Genes upregulated in high passage MSCs. FDR *q*-value = 0.0011. **(G)** Genes upregulated in elderly donor derived MSCs. FDR *q*-value = 0.0267. **(H)** Genes upregulated in senescent cells. FDR *q*-value = 0.0151.

**SI Figure 8.**
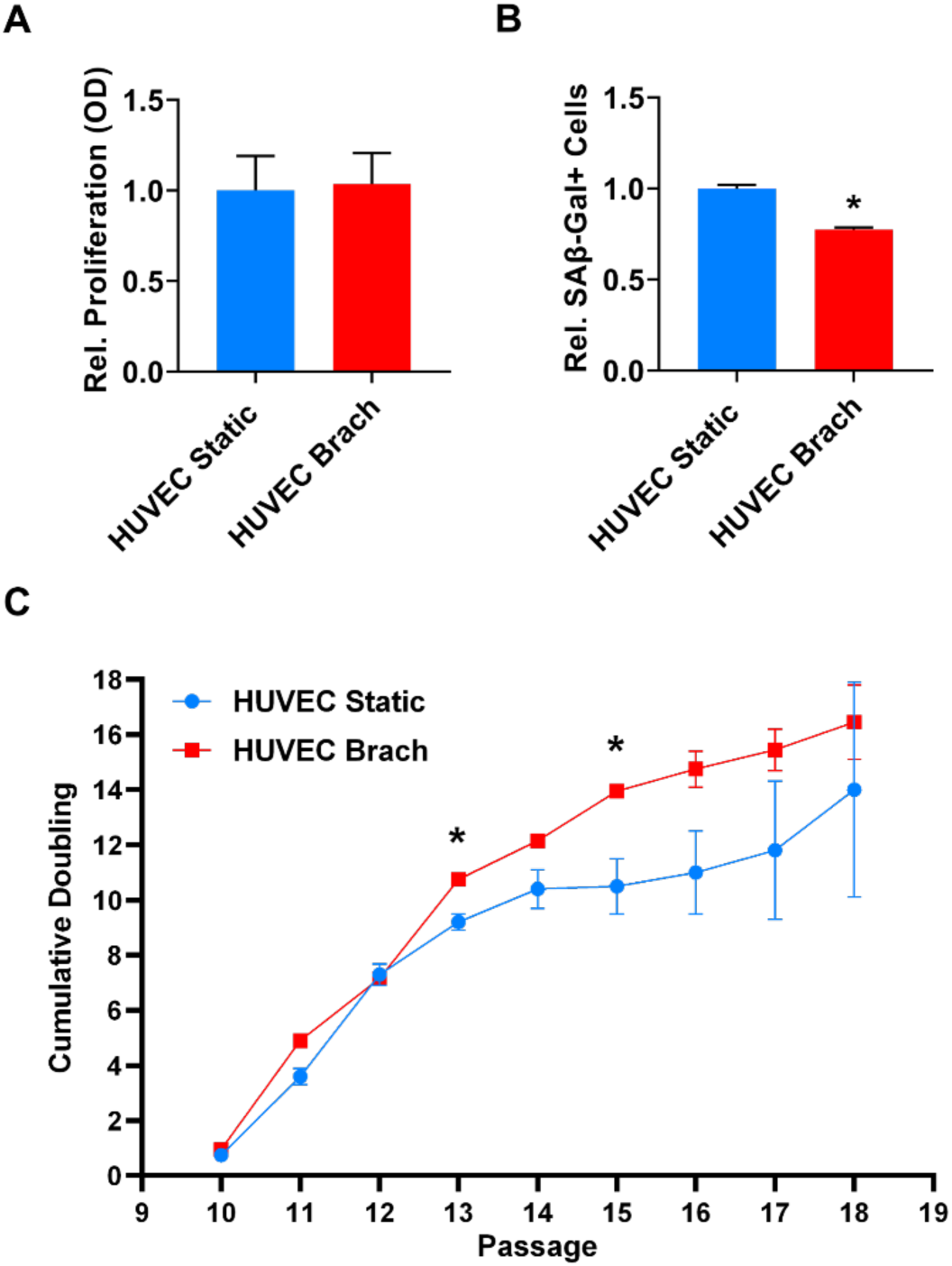
Human umbilical vein endothelial cells (HUVECs) at passage 10 were treated with brachial loading for 7 days and assessed for proliferative ability and functional phenotype. **(A)** Quantification of BrDU cell proliferation assay. **(B)** Quantification of fraction of β-galactosidase-positive cells. *p < 0.05 vs. static control treatment. **(C)** Quantification of the cumulative doubling of the treated cells during long-term culture expansion. *p < 0.05 vs. static control treatment.

**Supplemental Table 1.** Primary Antibodies Used for Immunoblotting.

| **Target Protein** | **Company** | **Catalog #** | **Species/Isotype** | **Dilution Ratio** |
| --- | --- | --- | --- | --- |
| Cyclin-D1 | Cell Signaling | 2978S | anti-rabbit | 1:500 |
| Phospho-Cyclin-D1<br>(Thr286) | Cell Signaling | 3300S | anti-rabbit | 1:500 |
| Akt | Cell Signaling | 4685S | anti-rabbit | 1:500 |
| Phospho-Akt (Ser473) | Cell Signaling | 9271S | anti-rabbit | 1:500 |
| SIRT1 | Cell Signaling | 9475S | anti-rabbit | 1:500 |
| SIRT3 | Cell Signaling | 2627S | anti-rabbit | 1:500 |
| Phospho-SIRT1<br>(Ser47) | Cell Signaling | 9787T | anti-rabbit | 1:500 |
| SIRT3 | Cell Signaling | 2627S | anti-rabbit | 1:500 |
| SIRT2 | Cell Signaling | 9787T | anti-rabbit | 1:500 |
| SIRT6 | Cell Signaling | 12486S | anti-rabbit | 1:500 |
| SIRT7 | Cell Signaling | 9787T | anti-rabbit | 1:500 |
| FOXO1 | Cell Signaling | 9946T | anti-rabbit | 1:500 |
| FOXO3a | Cell Signaling | 9946T | anti-rabbit | 1:500 |
| Phospho-FOXO4<br>(Thr28) | Cell Signaling | 9946T | anti-rabbit | 1:500 |
| Phospho-FOXO1(Ser256) | Cell Signaling | 9946T | anti-rabbit | 1:500 |
| Phospho-FOXO3a<br>(Ser253) | Cell Signaling | 9946T | anti-rabbit | 1:500 |
| SOD1 | Cell Signaling | 2770S | anti-rabbit | 1:500 |
| KU80 | Cell Signaling | 2180S | anti-rabbit | 1:500 |
| ATM | Cell Signaling | 2873S | anti-rabbit | 1:500 |
| Phospho-ATM<br>(Ser1981) | Cell Signaling | 5883S | anti-rabbit | 1:500 |

**Supplemental Table 2.** Primary Antibodies Used for Immunostaining.

| Target Protein | Company | Catalog # | Species/Isotype | Dilution Ratio |
| --- | --- | --- | --- | --- |
| FABP4 | R&D Systems | SC006 | anti-mouse | 1:500 |
| Osteocalcin | R&D Systems | SC006 | anti-mouse | 1:500 |
| Phospho-γ-H2AX<br>(Ser139) | Cell Signaling | 80312S | anti-mouse | 1:100 |
| KU80 | Cell Signaling | 2180S | anti-rabbit | 1:400 |

**Supplemental Table 3.** Reagents for Treating Cells During Mechanistic Studies.

| **Biomolecule** | **Company** | **Catalog #** | **Concentration** |
| --- | --- | --- | --- |
| EGFR-ErbB2-4<br>Inhibitor | Sigma-Aldrich | 324840 | 1 μM |
| Hydrogen Peroxide | Sigma-Aldrich | 216763 | 0.4 mM |
| N-Acetyl-L-Cysteine | Sigma-Aldrich | A9165 | 5 mM |
| KU-55933 | Selleck Chemicals | S1092 | 5 μM |
| Importazole | Cell Signaling | 10451S | 40 μM |
| Dooku 1 | Tocris | 6568 | 2 μM |
| GsMTx4 | Tocris | 4912 | 3 μM |

**Supplemental Table 4.** List of Primary Antibodies Used for Flow Cytometry.

| **Target Protein** | **Company** | **Catalog #** | **Fluor. Dye** |
| --- | --- | --- | --- |
| CD90 | R&D Systems | 967542 | APC |
| CD73 | R&D Systems | 967544 | Carboxyfluorescein |
| CD105 | R&D Systems | 967546 | PerCp/Cy5.5 |
| CD45 (Neg. Cocktail) | R&D Systems | 967548 | PE |
| CD34 (Neg. Cocktail) | R&D Systems | 967548 | PE |
| CD11b (Neg. Cocktail) | R&D Systems | 967548 | PE |
| CD79A (Neg. Cocktail) | R&D Systems | 967548 | PE |
| HLA-DR (Neg. Cocktail) | R&D Systems | 967548 | PE |

**Supplemental Table 5.** Gene Sets for GSEA Analysis.

| **Cell Type** | **GSE ID** |
| --- | --- |
| Human bone-marrow<br>mesenchymal stem cells | E-MTAB-4879 |
| Human bone-marrow<br>mesenchymal stem cells | GSE137186 |
| Human mixed cell types | CellAge (HAGR) |
| Human mixed cell types | Gene Expression Signature of Cellular<br>Senescence (HAGR) |
| Human mixed cell types | M27188 |

